# Trophic niche of the invasive gregarious species *Crepidula fornicata*, in relation to ontogenic changes

**DOI:** 10.1101/2020.07.30.229021

**Authors:** Thibault Androuin, Stanislas F. Dubois, Cédric Hubas, Gwendoline Lefebvre, Fabienne Le Grand, Gauthier Schaal, Antoine Carlier

**Affiliations:** ISMER, UQAR, Rimouski, Canada; IFREMER, DYNECO-LEBCO, Technopole Brest-Iroise, CS10070, Plouzané, France; UMR BOREA, Muséum National d’Histoire Naturelle, Concarneau, France; UMR CNRS 6539, LEMAR-IUEM-UBO, Plouzané, France

**Keywords:** *Crepidula fornicata*, trophic niche, ontogenic shift, fatty acids, stable isotopes, pigments, Bay of Brest

## Abstract

The slipper limpet *Crepidula fornicata* is a common and widespread invasive gregarious species along the European coast. Among its life-history traits, well documented ontogenic changes in behavior (i.e., motile male to sessile female) suggests a potential shift in feeding strategy across its life stages. Considering the ecological significance of this species in colonized areas, understanding how conspecifics share the trophic resource is crucial. Using fatty acids (FA) and stable isotopes (SI) as complementary trophic markers, we conducted a field survey between late winter and spring to investigate the trophic niche of three ontogenic stages of *C. fornicata* that bear different sexual (male/female) and motility (motile/sessile) traits. Potential trophic sources were characterized by their pigment, FA and SI compositions and discriminated well over the study period. We showed that the biofilm covering *C. fornicata* shells harbored a higher biomass of primary producers (i.e., chlorophytes and diatoms) than the surrounding sediment surface. Over the study period, we observed a covariation between the three ontogenic stages for both FA and SI compositions which suggest that the trophic niche of *C. fornicata* does not change significantly across its benthic life. During periods of low food availability, slipper limpets displayed an opportunistic suspension-feeding behaviour, relying on both fresh and detrital organic matter, likely coming from superficial sedimentary organic matter. During high food availability (i.e., spring phytoplankton bloom), all ontogenic stages largely benefited from this fresh supply of organic matter (pelagic diatoms in this case). However, the three ontogenic stages showed consistent differences in FA composition, and to a lesser extent in SI composition. These differences persist over time, as they originate more likely from ontogenic physiological changes (e.g., differential growth rates, metabolic rate or gametogenesis) rather than diet discrepancies.

## Introduction

The slipper limpet *Crepidula fornicata* is a non-indigenous and invasive gastropod originating from the East coast of the US (Hoagland, 1985). This species lives in the infralittoral zone but can be found from the intertidal zone down to 60 m depth. It has extensively colonized shallow soft bottom habitats of European coasts, from Norway to the Mediterranean Sea (Blanchard, 1997). Because of its introduction in many parts of the world and its potential cascading effect on food web functioning (Arbach Leloup et al., 2008; Chauvaud et al., 2000; Cugier et al., 2010), several studies have closely investigated its diet and inferred potential trophic overlap with co-occurring benthic species (Blanchard et al., 2008; Decottignies et al., 2007;Decottignies et al., 2007; Lefebvre et al., 2009; Riera, 2007; Riera et al., 2002). *C. fornicata* was overall considered as an opportunistic suspension-feeder, able to feed on a large array of trophic sources (e.g., phytoplankton, microphytobenthos, macroalgae, bacteria), depending on their availability. Based on stable isotope ratios, it has been hypothesised that there are a potentially large contribution of microphytobenthos, and more specifically benthic diatoms, in the diet of *C. fornicata* (Decottignies et al., 2007; Guérin, 2004; Lefebvre et al., 2009; Riera, 2007). However, the unexpected presence of inorganic carbonates in *C. fornicata* soft tissues have led to overestimate δ^13^C ratios in this consumer and then to overestimate the trophic role of microphytobenthos (Androuin et al., 2019).

*C. fornicata* is a protandrous hermaphrodites gregarious species. Sessile adults form stacks of several non-moving individuals, while juveniles and small males (~10 mm) are motile (Coe, 1936). Adult females are suspension-feeders, but contrary to bivalves, they lack labial palps and show no anatomical or functional potential for qualitative selection (Beninger et al., 2007). They form a food cord in a groove at the distal end of their gill filaments and ultimately catch their food with this cord with their radula before ingesting it (Shumway et al., 2014). For the related species *Crepidula fecunda*, which exhibits ontogenic behavior changes comparable to those of *C. fornicata*, it has been demonstrated that newly settled individuals first adopt a grazing feeding mechanism and gradually shift to a suspension-feeding behaviour once their gill is fully developed (Montiel et al., 2005). Young individuals are able to use both feeding mechanisms (i.e., grazing and suspension-feeding) during the motile phase of their life cycle (size < 28 mm), whereas females are exclusive suspension-feeders (Chaparro et al., 2002; Navarro and Chaparro, 2002). Such a shifting feeding mode has also been suggested for *C. fornicata* albeit without further behavioral evidence nor quantitative measurements. Using stomach content analysis, isolated adults of *C. fornicata* have been found to graze their surrounding habitat (Breton and Huriez, 2010). Younger individuals were also observed near grazing tracks but not analysed. By comparing the radula to body length ratio, Yee and Padilla (2015) found a more important role of the radula in small individuals (< 12.5 mm) and suggest a potential grazing behavior for those individuals. So, the relative contributions of the radula to feeding in different life stages of *C. fornicata* remain unclear (Shumway et al., 2014).

Since *C. fornicata* often occurs in large densities (up to 2000 ind. m^−2^) on the seafloor with all ontogenic stages grouped in stacks (Guérin, 2004; Martin et al., 2006), one can expect strong intraspecific interactions for food. These interactions could be either facilitative or competitive depending on ontogenic feeding ecology. While purely suspension-feeding slipper limpets should compete for food among ontogenic stages, recent works suggested that younger individuals may be facilitated by adults, both via a higher substrate availability (de Montaudouin and Accolla, 2018) and through the grazing of microphytobenthic biofilm colonizing adult shells (Androuin et al., 2018). In winter, when food is less available in the bay of Brest for suspension-feeders (Lavaud et al., 2018; Chatterjee et al., 2013), grazing may help young slipper limpets to avoid intraspecific competition for food. Moreover, *C. fornicata* often proliferates on muddy and turbid habitats with high suspended inorganic load, thus grazing behavior could also prevent the overloading of their digestive tract with inert matter of low nutritional quality (Navarro and Chaparro, 2002). Investigating the relationship (facilitation *vs*. competition) between age classes in a fecund invasive species like *C. fornicata* is crucial to better understand its population dynamic and so the consequences for the surrounding habitat.

Different trophic markers have long been used to investigate the trophic niche of marine benthic invertebrates (e.g., Blanchet-Aurigny et al., 2015; Cresson et al., 2016; Dubois and Colombo, 2014) and to describe the origin of assimilated particulate organic matter (hereafter OM) (Ke et al., 2017; Lavaud et al., 2018; Liénart et al., 2017). Carbon and nitrogen stable isotopes (SI) are broadly used to infer trophic niche of consumers (Fry and Sherr, 1984; Layman et al., 2012). Classically, nitrogen isotope ratio informs about the trophic position of a species and carbon isotope ratio reflects the origin of assimilated food sources (e.g., continental *vs*. oceanic, benthic *vs*. planktonic). In coastal ecosystems, the diet of most of benthic primary consumers is composed of a mixture of OM from various origins (phytoplankton, macroalgae, continental detritus, zooplankton, etc) which are often difficult to disentangle with isotopes of only two elements, namely carbon and nitrogen. This diversity of food sources implies that complementary trophic markers are relevant to complement SI informations (Majdi et al., 2018). For instance, pigment analyses have been widely used to study community composition of microscopic primary producers in the water column or in the sediment, since some pigments are specific to clades of algae (Brotas and Plante-Cuny, 2003; Roy et al., 2011). To a lesser extent, fatty acid (FA) compositions can be also specific of group of organisms, such as diatoms, bacteria, copepods or vascular plants (Dalsgaard et al., 2003; Kelly and Scheibling, 2012). FA represents the building stock of most lipid forms. They are energy reserves (neutral lipids) but also key structural component of cell membranes (polar lipids). In molluscs as for *C. fornicata*, such lipids storages are essential for larval development and during mature ontogenic stages for gonadal development (Deslous-Paoli and Héral, 1986; Leroy et al., 2013). Contrary to polar lipids, FA incorporated in the neutral fraction are directly mobilized and reflect more closely the FA composition of the diet (Fernández-Reiriz et al., 2015; Jezyk and Penicnak, 1966; Langdon and Waldock, 1981; Waldock and Nascimento, 1979). Extracting FA from this specific class of lipids and from a tissue with a rapid turnover (e.g., digestive gland, gonad) allows assessing rapid changes in the diet (McCutchan et al., 2003). Recently, the combined use of SI, FA and pigments improved our understanding of trophic pathways from the sources of particulate OM to benthic primary consumers (Lavaud et al., 2018; Majdi et al., 2018).

In this study, we investigated the extent of the trophic niche of *C. fornicata* and hypothesized that intraspecific differences exist in diet, associated with ontogenic behavior changes (i.e., motile male to sessile female). We expected that ontogenic trophic shift happens within stacks of *C. fornicata*, with a higher contribution of biofilm to motile males than to sessile males and females. To test our hypothesis, we conducted a field survey and characterized potential OM sources by their SI, FA and pigments compositions and inferred their assimilation in *C. fornicata* tissues using both SI and FA trophic markers.

## Methods

### Sampling strategy

The Bay of Brest (Brittany, France) is a 180 km^2^ semi-enclosed marine ecosystem. The sampling site is located near the Elorn estuary (48°23’N, 4°23’, average depth: 10 m) in a dense *C. fornicata* beds (~2000 ind. m^−2^) (Guérin, 2004) dominated by gravelly mud sediment (Gregoire et al., 2016). Potential OM sources and *C. fornicata* individuals were collected by SCUBA divers at five sampling dates (S1 = 26^th^ February, S2 = 21^th^ March, S3 = 28^th^ March, S4 = 12^th^ April and S5 = 14^th^ June) around mid and flood tide to ensure homogeneous mixing between estuarine and oceanic water. The late winter - spring period was chosen to encompass a period with potentially contrasting OM sources availability (e.g., spring blooms) (Figure 1).

**Figure 1:**
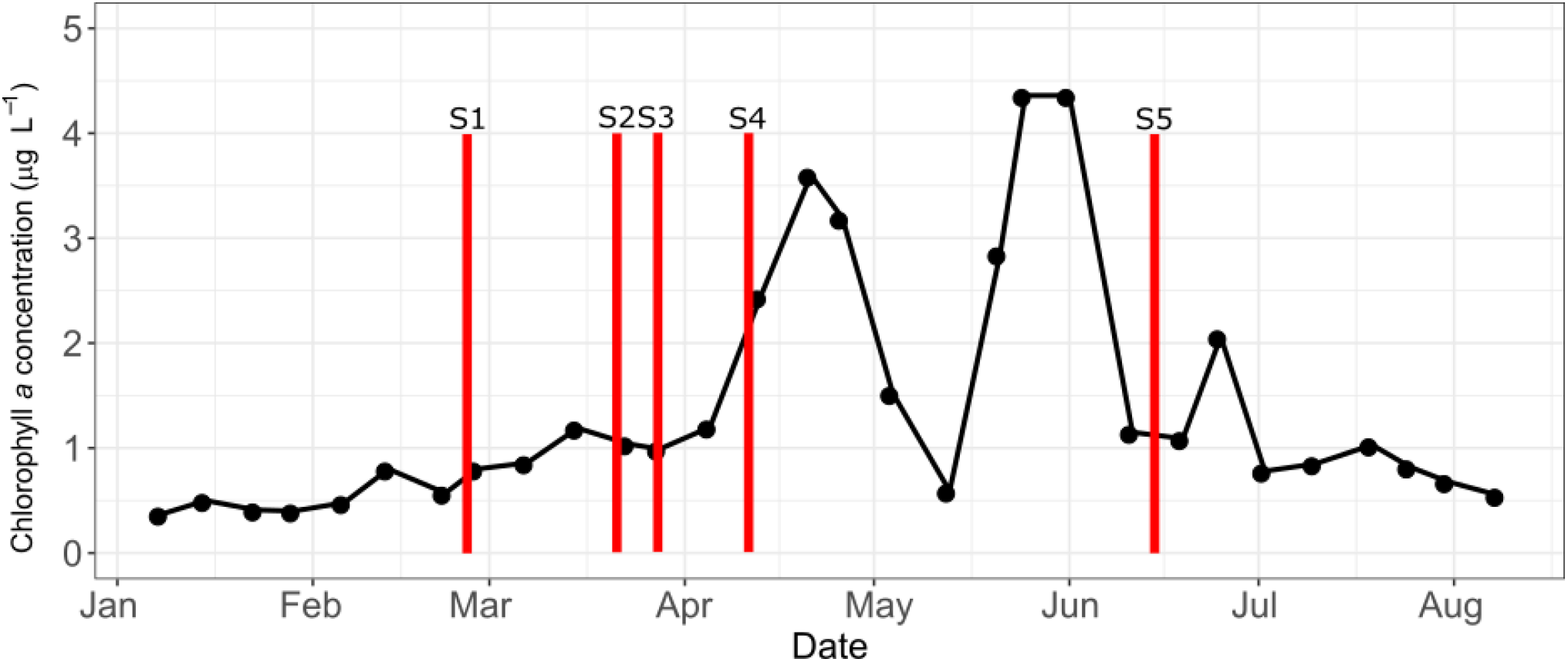
Sampling dates (S1 to S5) at the study site superimposed with weekly chlorophyll *a* concentration at the entrance of the bay of Brest in 2018 (data from the French Coastal Monitoring Network SOMLIT; http://somlit.epoc.u-bordeaux1.fr/fr/).

Suspended particulate organic matter (SPOM) was sampled at 50 cm above the sediment-water interface using two 8 L Niskin bottles, and immediately filtered on board onto a 200 μm nylon mesh to remove large zooplankton and particles. In the laboratory, between 1 and 1.5 L was filtered on pre-combusted (450°C for 5 hours) GF/F filters (0.7 μm). Three replicates for each of the three analyses (SI, FA and pigment) were obtained. Superficial sedimentary organic matter (SSOM) was sampled from three cores of 15 cm diameter and 15 cm depth. In the laboratory, the sediment-water interface was re-suspended by flushing seawater with a 30 mL syringe following a standardized process: 60 mL of SSOM was pre-filtered on a 200 μm nylon mesh to be consistent with SPOM samples and filtered on pre-combusted (450°C for 5 hours) GF/F filters (0.7 μm). Three replicates for each of the three analyses were obtained. Biofilm from one *C. fornicata* stack was scraped off using a toothbrush and suspended in 600 ml of filtered seawater (0.7 μm). 200 ml of the suspended solution was filtered on pre-combusted (450°C for 5 hours) on GF/F filters (0.7 μm). Three replicates for each of the three analyses were obtained. Filters for FA analysis were put in glass tubes containing 6 mL of chloroform-methanol (2:1, v:v) solution and stored at −80°C before analysis, whereas filters for pigment and SI analysis were immediately stored at −80°C.

Females of *C. fornicata* were sampled at the bottom of the stacks (mean shell length 33 ± 6 mm), attached to a dead *C. fornicata* shell. Sessile and motile males were sampled if they had a penis and a mean shell length of 20 ± 8 mm and 10 ± 1 mm, respectively. Smallest males were not necessarily found at the apex of the stack, fitting with the fact that they were fully motile and therefore potentially able to graze the substrate around them. We used the digestive gland as a relevant trophic integrator tissue because it has a higher turnover rate than muscle tissue and is an energy storage organ enriched in lipids (McCutchan et al., 2003; Vander Zanden et al., 2015). However, since digestive gland and gonad are fused in a single organ in *C. fornicata*, we analysed both tissues together for sessile males and females. Because gonads are comparatively small in motile males, the whole body was used to ensure sufficient lipid concentration. At each date and for each ontogenic stage, both stable isotope (SI) and fatty acid (FA) analyses of *C. fornicata* were performed on subsamples originating from the same tissue sample. Samples were stored at −80°C for a few days before freeze drying.

### Pigment analysis

The photosynthetic communities of SSOM, biofilm and SPOM have been analysed by the quantification of pigments by High Performance Liquid Chromatography (HPLC) according to Brotas and Plante-Cuny (2003). Filters were crushed and extracted in 3 ml of 95 % cold buffered methanol (2 % ammonium acetate) for 20 min at −20°C in the dark. Samples were centrifugated for 3 minutes at 3000 g after the extraction period. Extracts were then filtered with Whatman membrane filters (0.2 mm) immediately before HPLC analysis. Pigment extracts were analysed using an Agilent 1260 Infinity HPLC composed of a quaternary pump (VL 400 bar), a UV–VIS photodiode array detector (DAD 1260 VL, 190–950 nm), and a 100 μl sample manual injection loop (overfilled with 250 μl). Chromatographic separation was carried out using a C18 column for reverse phase chromatography (Supelcosil, 25 cm long, 4.6 mm inner diameter). The solvents used were A: 0.5 M ammonium acetate in methanol and water (85:15, v:v), B: acetonitrile and water (90:10, v:v), and C: 100 % ethyl acetate. The solvent gradient followed the Brotas and Plante-Cuny method (2003), with a flow rate of 0.5 mL min^−1^. Identification and calibration of the HPLC peaks were performed with chlorophyll *a*, ββ-carotene, chlorophyll c2, diatoxanthin, diadinoxanthin and fucoxanthin standards. All peaks detected were identified by their absorption spectra and relative retention times using the Open Lab CDS software (ChemStation Edition for LC/MS Systems, Agilent Technologies). Quantification was performed by repeated injections of standards over a range of dilutions to establish a standard curve of concentrations. Pigment percentages were expressed relatively to the surface/volume sampled (μg.cm^−2^ for biofilm and SSOM, and μg L^−1^ for SPOM). We measured the mean surface of three stacks of *C. fornicata* to standardize surfaces.

### Stable isotope analysis

δ^15^N and δ^13^C analyses were carried out independently for both OM sources and *C. fornicata* tissues. For OM sources, filters were freeze-dried and split in two equal parts. Half of the filter was scrapped off and weighed in tin capsules for δ^15^N analysis. The second half was decarbonated using acid-flume (10 N hydrochloric acid solution) for 7 hours (Lorrain et al., 2003), dried at 40 °C for 12 h, scrapped off and weighed in tin capsules for δ ^13^C analysis. *C. fornicata* samples were freeze-dried and ground into homogenous powder using a mortar and pestle. Approximately 400 μg of powder was weighed in tin capsules for δ^15^N analysis. Because both lipids content and inorganic carbonates can influence δ^13^C (Androuin et al., 2019; McCutchan et al., 2003), approximately 400 μg of powder was added to 1 ml of cyclohexane in Eppendorf tubes. Tubes were vortexed and centrifuged at 3000 g during 5 min. The supernatant was discarded, and the tubes dried at 40°C for a 12 h period. If the supernatant remained coloured, the sample was re-processed. Lipid-free tissues were then weighed in silver capsules and in-cup decarbonated using 1N HCl. Each capsule was visually checked, dried at 40°C for 1 h, and sealed. Samples were analysed for δ^15^N and δ^13^C by continuous flow on a Thermo Scientific Flash EA 2000 elemental analyser coupled to a Delta V Plus mass spectrometer at the Pôle de Spectrométrie Océan (PSO, Plouzané, France). Results are expressed in standard δ notation based on international standards (Vienna Pee Dee Belemnite for δ^13^C and atmospheric nitrogen for δ^15^N) following the equation:

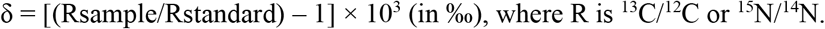

International isotopic standards of known δ^15^N and δ^13^C values were used: IAEA-600 Caffeine, IAEA-CH-6 Sucrose, IAEA-N-1 and IAEA-N-2 Ammonium Sulphate. The analytical precision was estimated using the standard deviation of an internal standard (Thermo Acetanilide, n = 8), as ± 0.11 ‰ and ± 0.07 ‰ for δ^13^C and δ^15^N values, respectively.

### Fatty acids analysis

Immediately after freeze-drying, tissues of *C. fornicata* were powdered and subsampled for FA analyses: between 2 and 20 mg - depending on ontogenic stages - were put in glass tubes (previously heated for 6 h at 450°C) containing 6 mL of a chloroform/methanol mixture (2:1, v:v), and extracted with a Dounce homogenizer. OM sources and *C. fornicata* samples were sonicated during 10 min and kept at −20°C until further analysis. The total lipid fractions were analysed in OM sources, whereas only the neutral lipids were analysed in *C. fornicata* samples. The detailed analysis method for separation and methylation is detailed in Le Grand et al. (2014). Fatty acid methyl esters (FAME) were analysed in a Varian CP 8400 gas chromatograph (GC) equipped with a split/splitless injector and a flame-ionization detector (FID). FAMEs were identified using two different capillary columns (ZBWAX 30 m × 0.25 mm i.d., 0.25 μm thickness, Phenomenex®; and ZB-5HT 30 m × 0.25 mm i.d., 0.25 μm thickness, Phenomenex®) by means of a standard FAME mix (S37, PUFA1, PUFA3, BAME, Sigma Aldrich®) and other lab-made standard mixtures with previously characterized and published FA compositions (e.g., non-methylene interrupted FA, Kraffe et al., 2004; Le Grand et al., 2013). FAs were expressed as the molar percentage of the total FA content.

### Statistical analyses

Pigment and FA compositions of OM sources, and FA compositions of ontogenic stages of *C. fornicata* were represented using a non-metric multidimensional Scaling (n-MDS). Homogeneity of the data was tested using permutational analyses of multivariate dispersion (PERMDISP) (Anderson, 2001). Statistical analyses on OM sources (pigments and FA) and *C. fornicata* (FA) were conducted using a non-parametric distanced-based permutation multivariate analysis of variance (PERMANOVA) based on a Bray-Curtis distance. Analyses were performed using two variables: OM sources or ontogenic stages (3 levels’ factors) and sampling dates (5 levels’ factor). Each date was considered independent due to the relative high turnover rate of both microorganisms found in the OM sources as well as the cells in the sampled tissues in *C. fornicata*. Following significant PERMANOVA results, *post hoc* tests were carried out using multiple pairwise comparisons with Bonferroni correction to identify differences among factors (Martinez Arbizu, 2017). However, the number of samples at each sampling date (3 < n < 5) was not sufficient to allow significant differences among the two factor levels in interaction, because of lack of statistical power when using Bonferroni correction in too many multiple comparisons. Therefore, *post hoc* comparisons of interaction term were not investigated. Finally, a SIMPER analysis was used to identify the FA explaining most of the dissimilarities between sampling dates and OM sources/ontogenic stages of *C. fornicata*.

Temporal variations and differences in SI ratios and pigment ratios/FA markers between OM sources/ontogenic stages of *C. fornicata* and sampling dates were assessed using two-way factorial analyses of variance (ANOVA). Because pigment and FA concentrations in SPOM were not comparable with biofilm and SSOM (surface *vs*. volume), SPOM was not include in the analysis. Its temporal variation was assessed using the non-parametric Kruskal-Wallis test. When ANOVA were significant, *post hoc* multiple comparisons were carried out using Tukey HSD. Normality and homogeneity of residuals were graphically assessed. When these conditions were not fulfilled, simple factor effect was verified using non-parametric Kruskal-Wallis and Mann-Whitney test for sampling dates and OM sources, respectively. Statistical analyses were performed in R version 3.3.0 (R Core Team, 2016) using packages ‘vegan’, ‘plyr’, ‘FactoMiner’, and ‘ggplot’.

## Results

### Organic matter sources

#### Pigments and fatty acids compositions

Overall, OM sources were well discriminated by their pigment compositions (Figure 2a), and by their FA compositions (Figure 2b).

**Figure 2:**
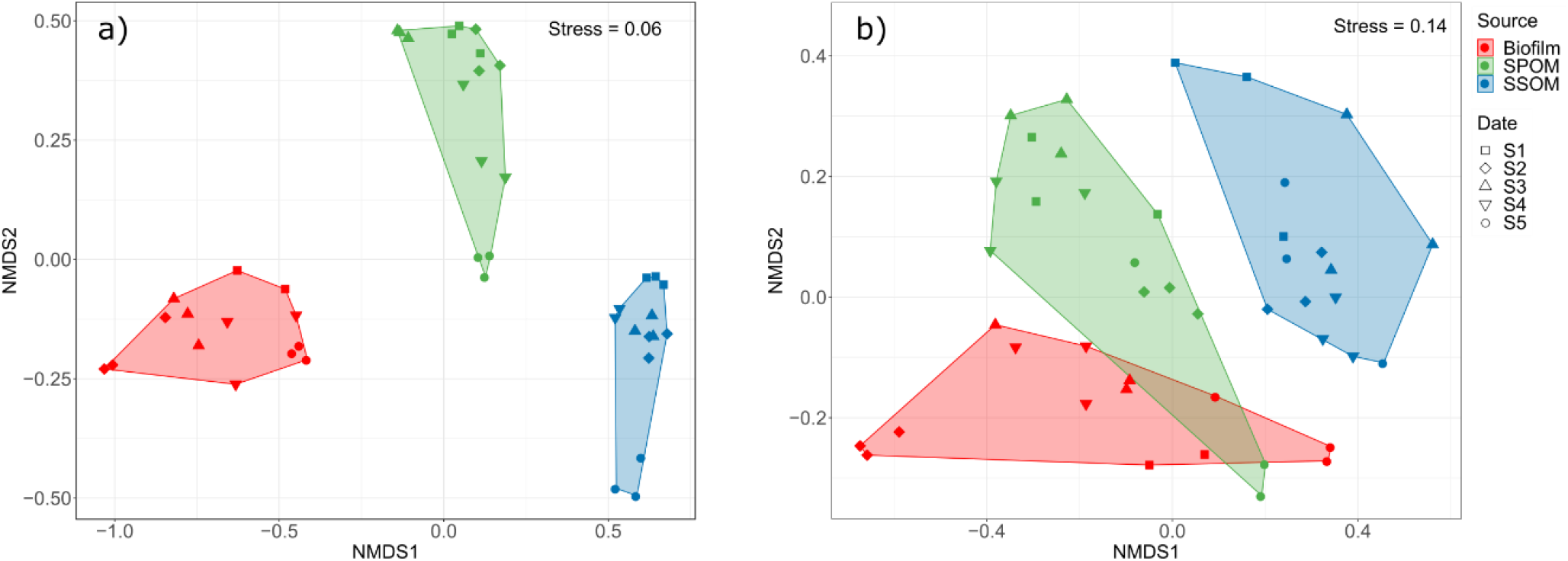
n-MDS based on the total pigment (a) and fatty acid (b) compositions of organic matter sources (Biofilm, suspended particulate organic matter (SPOM), superficial sedimentary organic matter (SSOM)). S1 to S5 correspond to the sampling dates (S1 = 26^th^ February, S2 = 21^st^ March, S3 = 28^th^ March, S4 = 12^th^ April and S5 = 14^th^ June).

Pigment compositions differed significantly between OM sources and sampling dates (*p* < 0.001 in both cases) and the interaction between the two factors was significant (*p* < 0.001). SIMPER analysis revealed that 80 % of the variability was explained by 7 pigments (Table S1). The biofilm was characterized by higher percentages of chlorophyll *b* and neoxanthin, together with one unknown pigment. SSOM was characterized by pheophytin *a*, pheophorbide *a*, and to a lesser extent lutein, whereas fucoxanthin and alloxanthin mainly discriminated SPOM. Temporal variations were mainly driven by a constant increase in both pheophorbide *a* and pheophytin *a* percentages for all OM sources. Fucoxanthin also showed an increase over time, except for SPOM and SSOM at S5 (Table S1).

Similarly, FA compositions also significantly differed between OM sources and sampling dates (*p* < 0.001 in both cases), and the two factors showed a significant interaction (*p* < 0.001). According to the SIMPER analysis, SSOM was characterized by higher percentages of 22:0, 16:1n-7, 18:1n-7 and a lower percentage of 16:0 (Table S2). The FA 18:0 and 20:4n-6 mostly discriminated biofilm whereas SPOM had higher percentages of 14:0 and 22:2n-6, and a lower percentage of 20:5n-3. In terms of temporal variations, both biofilm and SPOM showed similar decrease in saturated FA (i.e., 16:0 and 18:0) and increase in 16:1n-7 and 20:5n-3, especially between S4 and S5 (Table S2). SSOM exhibited less variable FA composition over time.

Total FA concentration did not show significant temporal variations for any OM sources (Figure 3a), with the biofilm always exhibiting higher concentration of total FA than SSOM over the sampling period (*p* < 0.001). Chlorophyll (chl) *a* concentration increased over time in PPOM up to sampling date S4 (1.8 ± 0.2 μg L^−1^) (*p* < 0.05) as well as in biofilm (reaching 3.9 ± 1 mg m^−2^ at S5) even if differences were not significant due to high between-samples variability (Figure 3b). Chl *a* concentration in SSOM slightly increased over time (*p* < 0.05) and was lower than in biofilm over the season (p < 0.001). Fucoxanthin concentration increased over time for all OM sources (*p* < 0.05 in all cases), followed by a decrease in S5 for SPOM and SSOM (Figure 3c). Fucoxanthin concentration was higher in biofilm than in SSOM (*p* < 0.001). The chl *b*: chl *a* ratio was 3 to 7-fold higher for biofilm than for SSOM (*p* < 0.001) and SPOM (*p* < 0.001), respectively (Figure 3d). Fucoxanthin: chl *a* (Figure 3e) and chl *c*: chl *a* (Figure 3f) ratios did not show clear temporal patterns for any OM sources and. However, they were 5 to 50-fold higher in SSOM than in SPOM (*p* < 0.001) and biofilm (*p* < 0.001) over the studied period, respectively.

**Figure 3:**
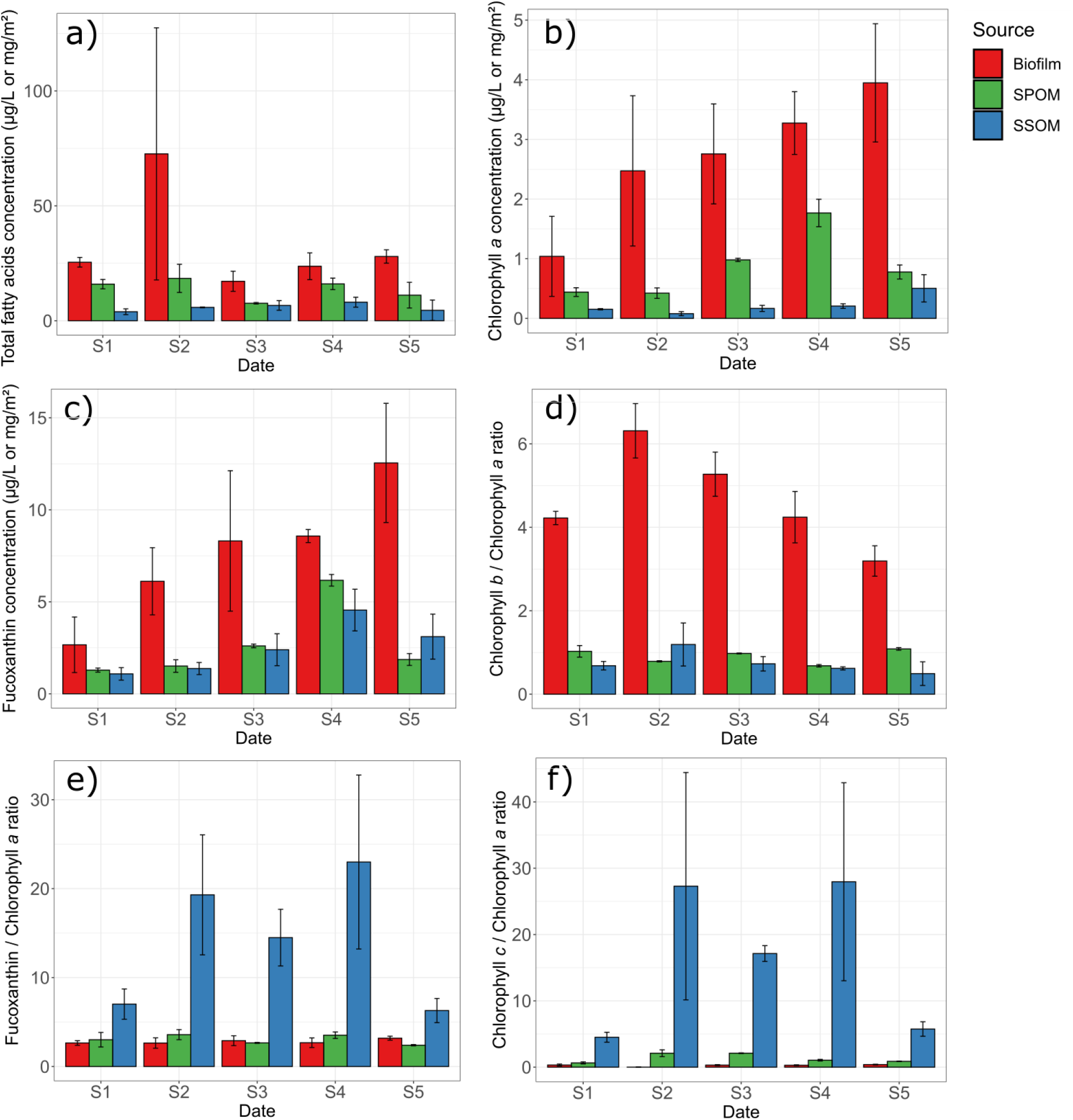
Total fatty acids (a), chlorophyll *a* (b) and fucoxanthin (c) concentrations, and ratios between chlorophyll *b* (d), fucoxanthin (e), and chlorophyll *c* (f) over chlorophyll *a* (mean ± SD, n ≥ 2) of organic matter sources (Biofilm, suspended particulate organic matter (SPOM), superficial sedimentary organic matter (SSOM)). S1 to S5 correspond to the sampling dates (S1 = 26^th^ February, S2 = 21^st^ March, S3 = 28^th^ March, S4 = 12^th^ April and S5 = 14^th^ June).

#### Stable isotopes composition

The three OM sources were well discriminated by their δ^13^C and δ^15^N values over the studied period. Their δ^13^C signal varied significantly according to both OM sources and sampling dates, and the interaction between the two factors was significant (*p* < 0.001, *p* < 0.01 and *p* < 0.001, respectively). PPOM was always depleted in ^13^C compared to RPOM (*p* < 0.001 at each date) and biofilm (*p* < 0.01 at each date) (Figure 4a, Table S3). Significant temporal δ^13^C variations were only observed in biofilm, with higher values at S5 than at S2 (*p* < 0.001) or S4 (*p* < 0.001). Biofilm was significantly enriched in ^15^N compared to both PPOM (*p* < 0.001) and RPOM (*p* < 0.001) (Figure 4b, Table S3). There was no interaction between OM sources and sampling dates (*p* = 0.26).

**Figure 4:**
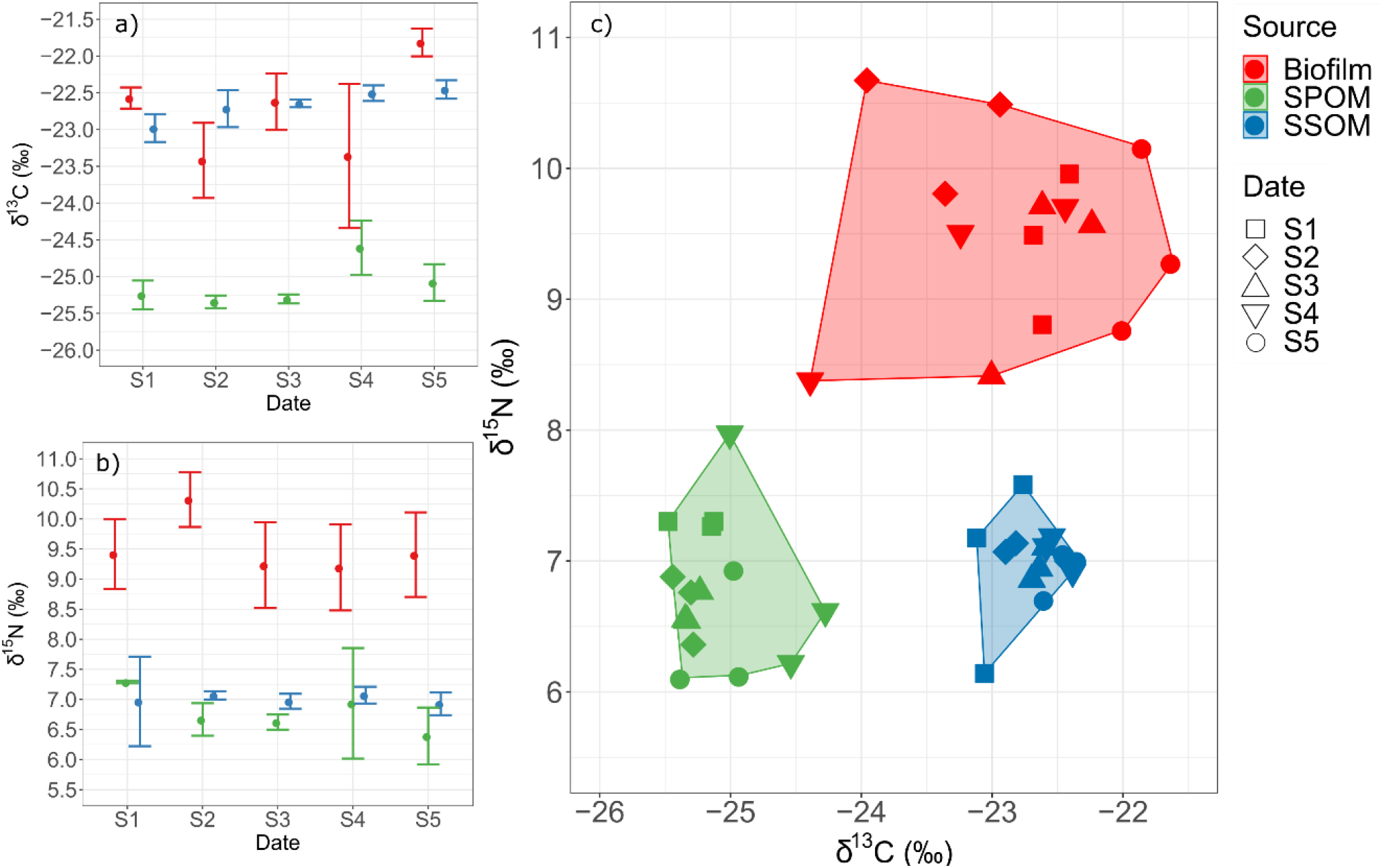
δ^15^N (a) and δ^13^C (b) isotopic compositions (mean ± SD, n = 3), and overall isotopic biplot (c) obtained of organic matter sources (Biofilm, suspended particulate organic matter (SPOM), superficial sedimentary organic matter (SSOM)). S1 to S5 correspond to the sampling dates (S1 = 26^th^ February, S2 = 21^st^ March, S3 = 28^th^ March, S4 = 12^th^ April and S5 = 14^th^ June).

### Crepidula fornicata

#### Fatty acids composition

FA composition of *C. fornicata* significantly differed between ontogenic stages and sampling dates (*p* < 0.001 in both cases) and the interaction between the two factors was significant (*p* < 0.001). The analysis of multivariate dispersion was also significant (*p* < 0.05), indicating that multivariate dispersion was not homogeneous. This was clearly illustrated by the n-MDS (when comparing the convex hull areas) where motile males showed much higher variation than sessile females and sessile males exhibited an intermediate level of variation (Figure 5). Pairwise SIMPER analyses between ontogenic stages revealed that sessile males were mainly characterized by saturated FA 16:0 and 18:0, especially at the two first sampling dates (Table S4). Sessile females differed from both motile and sessile males by higher percentages of C_20_ FA such as 20:5n-3, 22:6n-3 and 20:1n-11, but also higher percentages of odd branched FA (Table S4). Sessile males showed an overall comparable FA composition than sessile females but exhibited higher variability between sampling dates, as shown by the n-MDS. The SIMPER analyses performed between dates revealed that FA that most contributed to the observed temporal changes were the FA 16:0, 18:0 and 22:6n-3 decreasing over time, and the FA 20:5n-3 and 16:1n-7 increasing over time, all accounting for approximately 45 % of the dissimilarity over the 5 sampling dates (Table S4).

**Figure 5:**
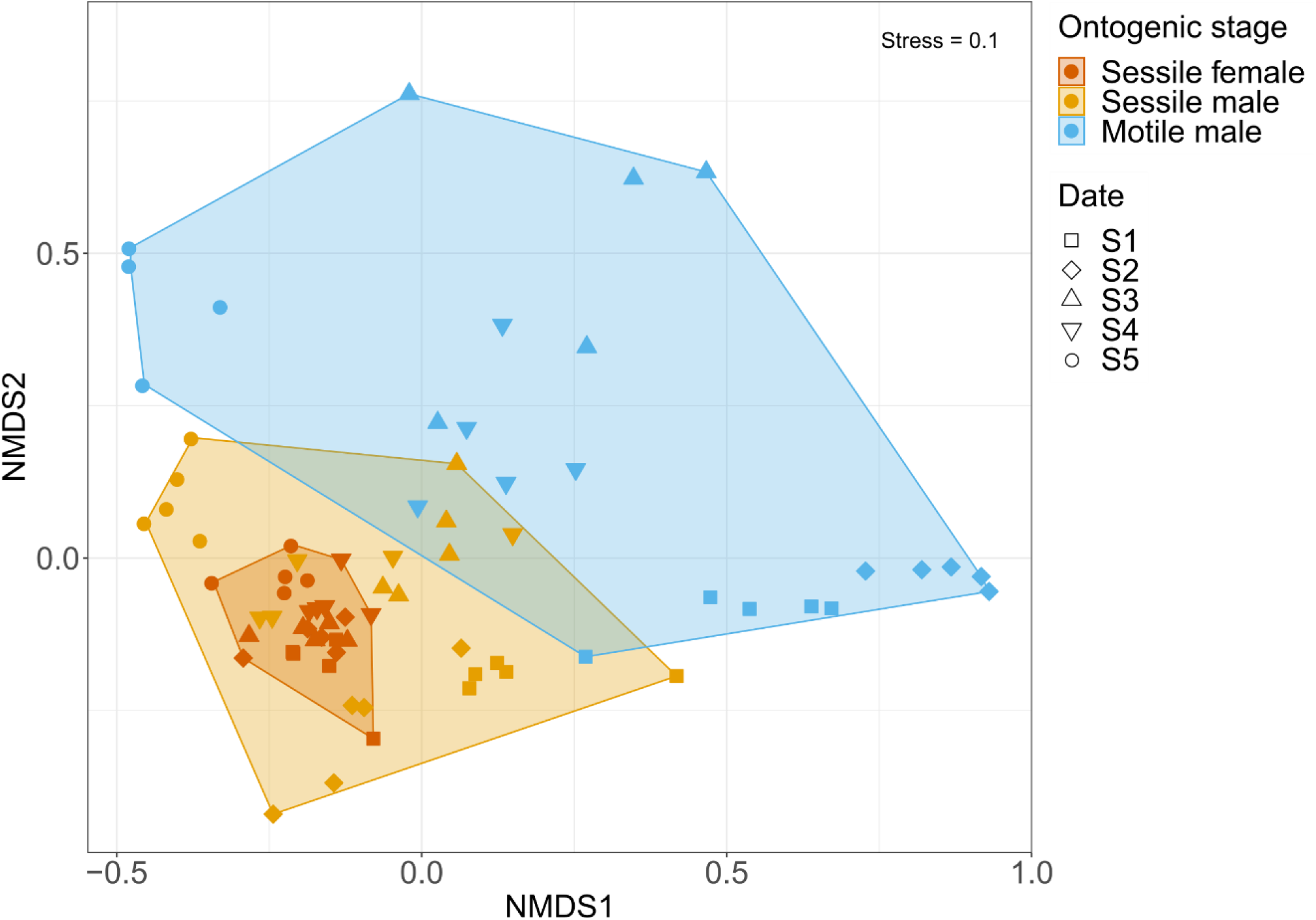
n-MDS based on the total fatty acid compositions of ontogenic stages of *Crepidula fornicata* (motile males, sessile males, sessile females). S1 to S5 correspond to the sampling dates (S1 = 26^th^ February, S2 = 21^st^ March, S3 = 28^th^ March, S4 = 12^th^ April and S5 = 14^th^ June).

All FA percentages and ratios changed significantly between sampling dates (*p* < 0.001 in all cases) and between ontogenic stages (*p* < 0.05 in all cases, except for 16:1n-7 and 18:4n-3). The interaction terms were always significant (*p* ≤ 0.05). Overall, temporal variations in FA composition were the highest in motile males, the lowest for females and intermediate for sessile males.

PUFA/SFA ratio increased over time for both motile and sessile males but remained constant in females (Figure 6a). There was no significant difference between ontogenic stages at sampling date S5. The relative abundance of branched FA was quite variable but significantly higher in sessile females than in motile males (*p* > 0.05) (Figure 6b). The highest values in branched FA was recorded in sessile females at S1 (11.2 ± 4.3) and the lowest in sessile males at S5 (3.8 ± 1). The ratio between 20:5n-3 and 22:6n-3 exhibited temporal variations for all ontogenic stages (Figure 6c), with a strong increase in S5 (*p* < 0.001 in all cases) where values ranged from 2.5 ± 0.5 in sessile females to 4 ± 0.6 in motile males. The FA 16:1n-7 followed the same trend as the 20:5n-3 / 22:6n-3 ratio with a more gradual increase over time for all ontogenic stages (Figure 6d). The highest values (from 3.6 ± 0.9 in sessile females to 6.4 ± 1.9 in motile males) were also recorded at S5. The FA 18:4n-3 also showed strong temporal variations in all ontogenic stages (Figure 6e), with an increase up to S4 followed by a decrease at S5. Finally, the n-3/n-6 ratio showed no temporal variation for sessile females but increased significantly over time up to S4 in both sessile (*p* < 0.01 in all cases) and motile males (*p* < 0.05 in all cases) (Figure 6f).

**Figure 6:**
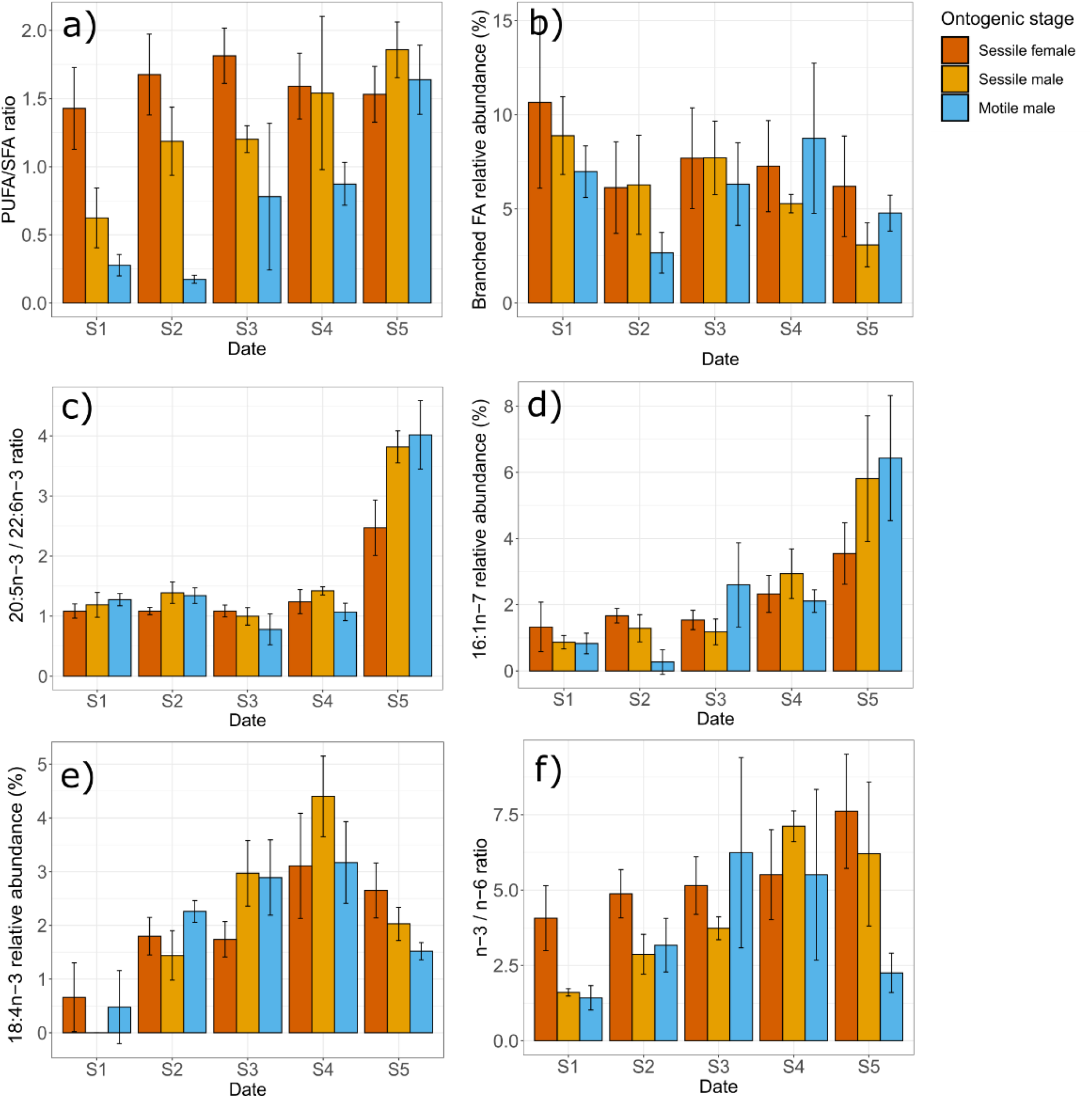
Relative abundance of fatty acids (FA) in the three ontogenic stages of *Crepidula fornicata* (motile males, sessile males, sessile females) (mean ± SD, n = 5): (a) Polyunsaturated FA / Saturated FA ratio, (b) Branched FA, (c) 20:5n-3 / 22:6n-3 ratio, (d) 16:1n-7, (e) 18:4n-3 and (f) n-3/n-6 ratio. S1 to S5 correspond to the sampling dates (S1 = 26^th^ February, S2 = 21^st^ March, S3 = 28^th^ March, S4 = 12^th^ April and S5 = 14^th^ June).

#### Stable isotopes composition

Overall, the three ontogenic stages exhibited similar isotopic patterns over time (Figures 7a and 7b, respectively). No interactions were found between ontogenic stage and sampling date, both for carbon (*p* = 0.6) and nitrogen (*p* = 0.27). Motile males were significantly depleted in ^13^C compared to sessile females (*p* < 0.05) and significantly depleted in ^15^N compared to both sessile females (*p* < 0.001) and males (*p* < 0.05). Significant temporal variations were observed for δ^13^C (*p* < 0.001) with a marked ^13^C enrichment at S5 compared to all other sampling dates (*p* < 0.05 in all cases), up to 2 ‰ when compared to S4. A significant temporal decrease in δ^15^N was found (*p* < 0.001) for all ontogenic stages.

**Figure 7:**
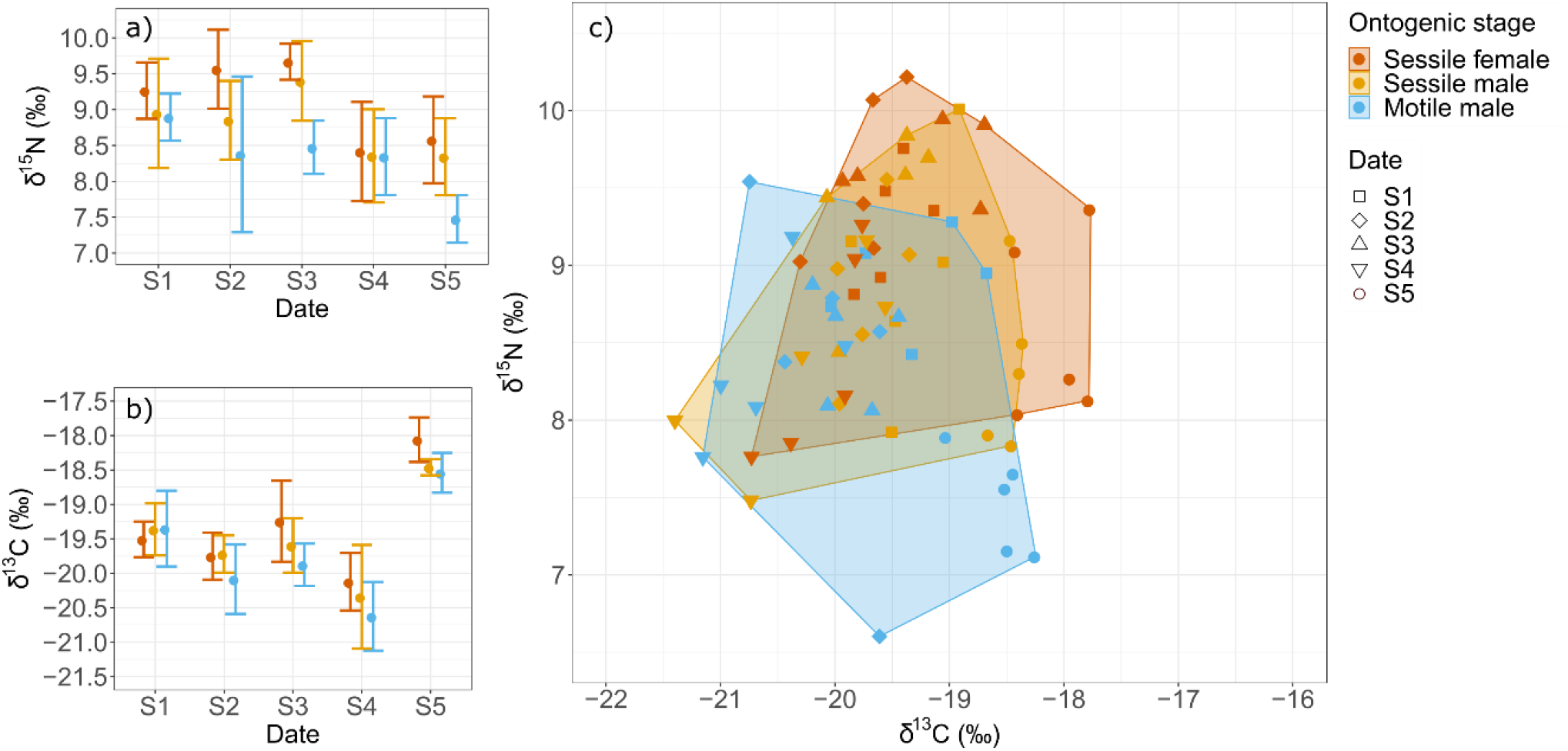
δ^15^N (a) and δ^13^C (b) isotopic compositions (mean ± SD, n = 5), and corresponding isotopic biplot (c) for three ontogenic stages of *Crepidula fornicata* (motile males, sessile males, sessile females). S1 to S5 correspond to the sampling dates (S1 = 26^th^ February, S2 = 21^st^ March, S3 = 28^th^ March, S4 = 12^th^ April and S5 = 14^th^ June).

## Discussion

Contrary to our hypothesis, and while the OM sources were well discriminated by their pigments, FA and SI compositions, trophic markers measured in *Crepidula fornicata* suggested an overall similar trophic niche across its ontogenic stages. Our results confirmed that the slipper limpet is an opportunistic suspension-feeder that exploits both pelagic and benthic particulate OM in varying proportions according to the season and sources availability. Although differences in FA composition, and to a lesser extent in SI composition, were noticeable between ontogenic stages at each sampling date, we think they likely reflect ontogenic physiological changes linked to differential growth rate and energetic demand rather than profound changes in diet.

### Composition and availability of potential food sources

δ^13^C in SPOM (−25.4 ± 0.7 ‰) in our study site was likely influenced by both marine (−22.3 ± 1.2 ‰, data from the French Coastal Monitoring Network SOMLIT; http://somlit.epoc.u-bordeaux1.fr/fr/) and terrestrial POM (−27.5 ± 1.2 ‰, Mortillaro et al., 2014), as mentioned by Marchais et al. (2013). Over our study period, SPOM showed comparable chlorophyll *a* biomass (0.4 - 1.8 μg L^−1^) than currently encountered in other areas of the bay of Brest (0.3 - 5 μg L^−1^, Chatterjee et al. 2013). High percentages of fucoxanthin and alloxanthin suggested the presence of Bacillaryophyta (i.e., diatoms) and Cryptophyta in the water column, respectively (Brotas and Plante-Cuny, 2003; Roy et al., 2011). Using the SOMLIT weekly monitoring of chlorophyll *a* and marine POM δ^13^C, a typical ^13^C enrichment (~3 ‰) was noticeable during a phytoplankton bloom that occurred between May 25^th^ and June 1^st^ in the bay. This bloom was mainly composed by three diatoms: *Cerataulina pelagica* (1.1 10^6^ cells L^−1^), *Leptocylindricus danicus* (4.7 10^4^ cells L^−1^) and *Rhizosolenia imbricata* (1.2 10^4^ cells L^−1^) (data extracted from the REPHY network, IFREMER). While this bloom was not sampled in the PPOM sampling set, the ^13^C enrichment was reflected in the tissues of *C. fornicata* collected at S5 (14^th^ June).

Among the three sources of organic matter, SSOM showed the most homogeneous isotopic composition over time, which is unexpected as SSOM is often considered as a mixture of pelagic and benthic OM sources and consequently highly variable according to OM sources proportions and isotopic compositions. SSOM appeared here as a complex mixture of low and high quality OM (Lefebvre et al., 2009; Rigolet et al., 2014). On one hand, it was characterized by i) pheophorbide *a* and pheophytin *a*, which are degradation products of chlorophyllide *a* and chlorophyll *a*, respectively (Brotas and Plante-Cuny, 1998; Cartaxana et al., 2003), ii) odd branched FA (such as ant15:0) and the 18:1n-7 indicating the presence of bacteria (Hubas et al., 2017; Jaschinski et al., 2011; Meziane et al., 1997) and iii) the long chain saturated fatty acids 22:0 which confirms the presence of a refractory terrestrial contribution in this environment (Canuel, 2001). On the other hand, SSOM exhibited the highest PUFA/SFA ratio most of the time (Table S1), suggesting higher quality/lability compared to SPOM and biofilm (Connelly et al., 2015, 2016; Parrish et al., 2005). This was confirmed by high Fucoxanthin / Chl *a* and Chl *c* / Chl *a* ratios which indicated a higher contribution of diatoms in SSOM than in SPOM and biofilm (Brotas and Plante-Cuny, 2003).

Biofilm scraped on shells of *C. fornicata* showed higher chlorophyll *a* concentration than the surrounding sediment, suggesting higher biomass of primary producers on shells (Androuin et al., 2018). The high percentages of chl *b* and neoxanthin, as well as the Chl *b* / Chl *a* ratio in the biofilm also suggest that chlorophytes were abundant on shells (Brotas and Plante-Cuny, 2003). While these results were not supported by FA (e.g., 16:4n-3, 18:3n-3 or 18:4n-3 characterizing chlorophytes, Fleurence et al. 1994; Kelly and Scheibling, 2012), mollusc shells are known to harbour microchlorophytes or macrochlorophytes propagules (Barillé et al., 2017; Mineur et al., 2007). FA characterizing the biofilm were 18:0 and 20:4n-6. While 18:0 is an ubiquitous FA in marine environment, the 20:4n-6 can be found in large proportion in red and brown algae (Fleurence et al., 1994; Kelly and Scheibling, 2012). It is worth noting that *C. fornicata* shells were partly covered with crustose red algae in our study site (pers. obs.). The high concentration of fucoxanthin also suggests the presence of high biomass of diatoms on these shells, as already mentioned by Ní Longphuirt et al. (2007).

### Trophic niche of *C. fornicata*, in relation to ontogenic changes

Overall, SI ratios of *C. fornicata* showed that the three ontogenic stages had similar isotopic niches, although the niche of sessile females does not fully overlap with those of motile males. According to the respective SI ratios of potential food sources (Biofilm, SPOM, SSOM and marine POM) and those of *C. fornicata* tissues, and considering classical diet to consumer trophic enrichment factor (~ 0.75 - 1 ‰ for carbon and ~2.5 - 2.74 ‰ for nitrogen, (Caut et al., 2009; McCutchan et al., 2003), it is likely that *C. fornicata* relies either on SSOM or marine POM depending of the season and food availability in the water column. During lower food availability period (i.e., end of winter and early spring), it is difficult to disentangle SSOM from marine POM since their SI signals do not differ. However marine POM was sampled close to the Bay entrance with higher oceanic influence, whereas SPOM was sampled just above the *C. fornicata* beds around mid and flood tide to ensure mixing with oceanic water. Therefore, SPOM was more readily available than marine POM for the slipper limpet. It is then reasonable to assume that SI composition of *C. fornicata* refers to SSOM rather than marine POM, since SSOM can be exploited through regular resuspension events linked to tidal currents (Beudin, 2014). After the phytoplankton spring bloom that occurred in the bay of Brest at the end of May, producing a ^13^C enrichment in the water column (~3 ‰), a similar ^13^C enrichment was found for all stages of *C. fornicata* (~2 ‰). These results clearly showed that adults as well as young individuals of *C. fornicata* benefited from the spring bloom. However, minor but consistent isotopic differences were also found between ontogenic stages at each sampling date, which cannot necessarily be attributed to an ontogenic diet shift. Indeed, inferring diet shift using SI ratios may be hampered by the effects of physiological changes occurring during ontogeny such as gonadal maturation, metabolic rate or differential tissue growth between youngs and adults (Blanchet-Aurigny et al., 2012; Hentschel, 1998; Rossi et al., 2004; Vander Zanden et al., 2015). Lefebvre and Dubois (2016) analysed trophic enrichment factor and turnover rate in several marine benthic invertebrates, including *C. fornicata*. They showed a clear negative relationship between growth and enrichment factor values. Young motile males of *C. fornicata* are growing more rapidly than larger sessile males or even larger females (Hoagland, 1978; Walne, 1956). Enrichment factors are then expected to be smaller for motile males than sessile males or females. This is consistent with small between-stage differences found in SI ratios (Figure 7c). Even if motile males were analysed *in toto* (including muscle tissue with longer turnover rate), we believe that this does not bias interpretations of between-stages differences over the study period. Analysing only tissues with fast turnover rate (as we did for sessile individuals) would have increased these between-stages differences.

The FA profiles of the three ontogenic stages of *C. fornicata* showed a consistent temporal variation over the study period, suggesting similar changes in food sources utilization. This temporal pattern resulted from three distinct groups of sampling dates (Figure 5). The two first ones (S1 and S2) likely corresponded to a period that integrated the trophic signal of winter season’s food sources, whereas the last one (S5) clearly showed the assimilation of the spring phytoplankton bloom. The two-intermediate sampling dates (S3 and S4) corresponded to a transition reflecting an increase in food availability. As interpreted above with SI, all ontogenic stages of *C. fornicata* may have probably exploited SSOM before the spring bloom when food in the water column is less available. In this pool of OM, FA revealed that slipper limpets likely fed on benthic diatoms (as suggested by 16:1n-7 and 20:5n-3; Dunstan et al. 1992; Napolitano et al. 1997; Passarelli et al. 2012), dinoflagellates (22:6n-3; Zhukova and Aizdacher 1995; Lavaud et al. 2018) and bacteria (Branched FA and 18:1n-7; Perry et al. 1979; Zhukova et al. 1992; Haack et al. 1994), which is in agreement with previous interpretations done in other coastal bays (Dubois et al., 2014; Leroy et al., 2013). In sediment, bacteria are often associated with detritus and are therefore not considered as a high-quality food source (Dalsgaard et al., 2003). This is confirmed by the PUFA/SFA ratio, a biomarker of fresh *vs*. detritic OM (Connelly et al., 2015, 2016; Parrish et al., 2005), which was lower in the slipper limpet (~ 1.5, our study) than in other suspension-feeding species of the bay of Brest, such as *Pecten maximus* (~ 2.8, Lavaud et al. 2018) or *Ophiotrix fragilis* (~ 2, Blanchet-Aurigny et al. 2015). The fact that *C. fornicata* lacks pre-ingestive mechanisms for particle selection likely explains their opportunistic trophic behaviour based on both fresh and detritic organic matter (Beninger et al., 2007). After spring bloom, the percentages of diatom’s markers 16:1n-7 and 20:5n-3 drastically increased in *C. fornicata*’s tissue, as well as the 20:5n-3/22:6n-3 ratio, confirming that the three ontogenic stages benefit from this food supply from the water column (Budge and Parrish, 1998; Lavaud et al., 2018). Finally, the contribution of FA 18:4n-3 increased over time for all ontogenic stages. According to the literature, this FA may originate from different primary producers such as dinoflagellates (Budge and Parrish, 1998), brown or green macroalgae (Fleurence et al., 1994; Kelly and Scheibling, 2012). Considering i) the absence or low temporal variation of other dinoflagellate biomarkers (peridinin pigment and 22:6n-3 FA) in the OM sources, ii) the presence of brown algae on the rocky shore, and iii) the frequent seasonal accumulation of green macroalgae in our study area (Study Centre for Algal Promotion, http://www.ceva.fr; Ragueneau et al., 2018), we can expect a seasonal trophic role of these macroalgae for *C. fornicata* at our study site, probably in the form of detrital particles. FA dietary biomarkers in consumers are subject to physiological regulation (e.g., specific retention, *de novo* synthesis) during uptake and trophic transfer, and can be species-specific (Galloway and Budge, 2020). Although the use of neutral lipids theoretically limits this shortcoming, dedicated experimental studies on FA metabolism and FA trophic transfer in marine limpets specifically are still needed (Galloway and Budge, 2020; Zhukova, 2019).

Contrary to SI, FA compositions of the three ontogenic stages of *C. fornicata* showed low overlap, especially between motile and sessile limpets. Considering that lipids and fatty acids profiles may be age and sex-specific (Correia et al., 2003; Pernet et al., 2012), some of the differences in FA compositions are then likely to originate from physiological changes between ontogenic stages. During winter period, motile males were characterized by higher proportions of SFA such as 16:0 (25-34 %) and 18:0 (27-36 %). These FA are very common in marine organisms and do not necessarily reflect a specific diet (Dalsgaard et al., 2003; Kelly and Scheibling, 2012). Moreover, despite the dietary interest of short-chain SFA, where energy is more efficiently released via beta-oxidation than for PUFA (Langdon and Waldock, 1981), temperature may also influence the process of their utilization (Pernet et al., 2007). For example, it has been experimentally demonstrated that cold-acclimated oysters (5-7°C) have a clear preference for PUFA (n-3) over SFA (16:0) as fuel for energy compared to ‘temperate’ oysters (Chu and Greaves, 1991). The lower utilization of SFA in cold-acclimated oysters has been attributed to the fact that SFA are not in the liquid phase under cold temperature, thus making them less accessible for catabolic processes. In *Crepidula fornicata*, young individuals have proportionally less energy reserves than adults (Guérin, 2004) and are more subjected to low temperature effects due to a higher surface-to-volume ratio (Diederich et al., 2015). Hence, the lower utilization of SFA could explain their higher SFA percentages during the winter period in the bay of Brest, where temperatures fall down to 7°C (Figure S1). Moreover, the weight-specific metabolic rate, which is higher in smaller organisms (Bayne and Newell, 1983; Bougrier et al., 1995), could be exacerbated in *C. fornicata* because motile males are more active than adults through their motility (Coe, 1936; Hoagland, 1978; Walne, 1956). Together, this may explain the higher temporal and inter-individual variability in their FA compositions. It also suggests that young motile individuals of *C. fornicata*, having less energetic storage in winter while having more energetic needs, are probably in poor energetic condition during this period.

As we measured neutral FA in both digestive gland and gonad simultaneously (because the digestive gland cannot be isolated from the gonad), the level of lipid storage and FA composition may also depend on their sexual development stage. Indeed, females allocate more energy than males in the reproduction due to maternal gametogenesis (Deslous-Paoli and Héral, 1986; Leroy et al., 2013). As an illustration of FA composition changes, the n-3/n-6 ratio increased over time for both sessile and motile males whereas it remains unchanged in females. This may be linked to a preferential allocation of n-3 to early embryos, which showed an increase over the reproductive period of the slipper limpet (Leroy et al., 2013).

### Conclusion

Contrary to our hypothesis, *Crepidula fornicata* did not show ontogenic feeding shift, despite its peculiar way of life, and development from motile male to sessile female. However, we confirmed the opportunistic feeding behaviour of this suspension-feeder in our study site. Although we only found minor between-stage differences in both SI and FA compositions, it is likely that these differences originate from physiology rather than trophic behavior. These results suggest that young *C. fornicata* (i.e., one year old) may compete for food with adults. This intraspecific competition is expected to be higher in winter, when food is more limited and when young *C. fornicata* are probably in poorer condition. In a recent work, de Montaudouin and Accolla (2018) showed a recruitment facilitation through higher substrate availability in high density beds with a potential mitigation by the negative effect of intraspecific competition. Our study supports this idea of ontogenic trophic competition in *C. fornicata* when food is limiting.

## Acknowledgements

We thank the PSO and LIPIDOCEAN analytical facilities for stable isotope and fatty acid facilities, respectively (Oanez Lebeau, Antoine Bideau and Rudolph Corvaisier). We are grateful to Aline Blanchet-Aurigny for commenting upon preliminary versions of this manuscript. We also thank the LEBCO diving team (Amelia Curd, Xavier Caisey and Aurélien Tancray) for providing biological samples. We finally thank two anonymous reviewers for valuable input for the manuscript. TA was funded by an IFREMER, LabexMER and Region Bretagne PhD grant. This work was funded by the TOTAL foundation for biodiversity. Version 4 of this preprint has been peer-reviewed and recommended by Peer Community In Ecology (https://doi.org/10.24072/pci.ecology.100077).

## Data accessibility

Raw data and R-scripts are freely available at: https://www.seanoe.org/data/00646/75809/

## Conflict of interest disclosure

The authors of this preprint declare that they have no financial conflict of interest with the content of this article.

Cédric Hubas is one of the PCI Ecology recommenders.

## Supplementary materials

**Table S1:**
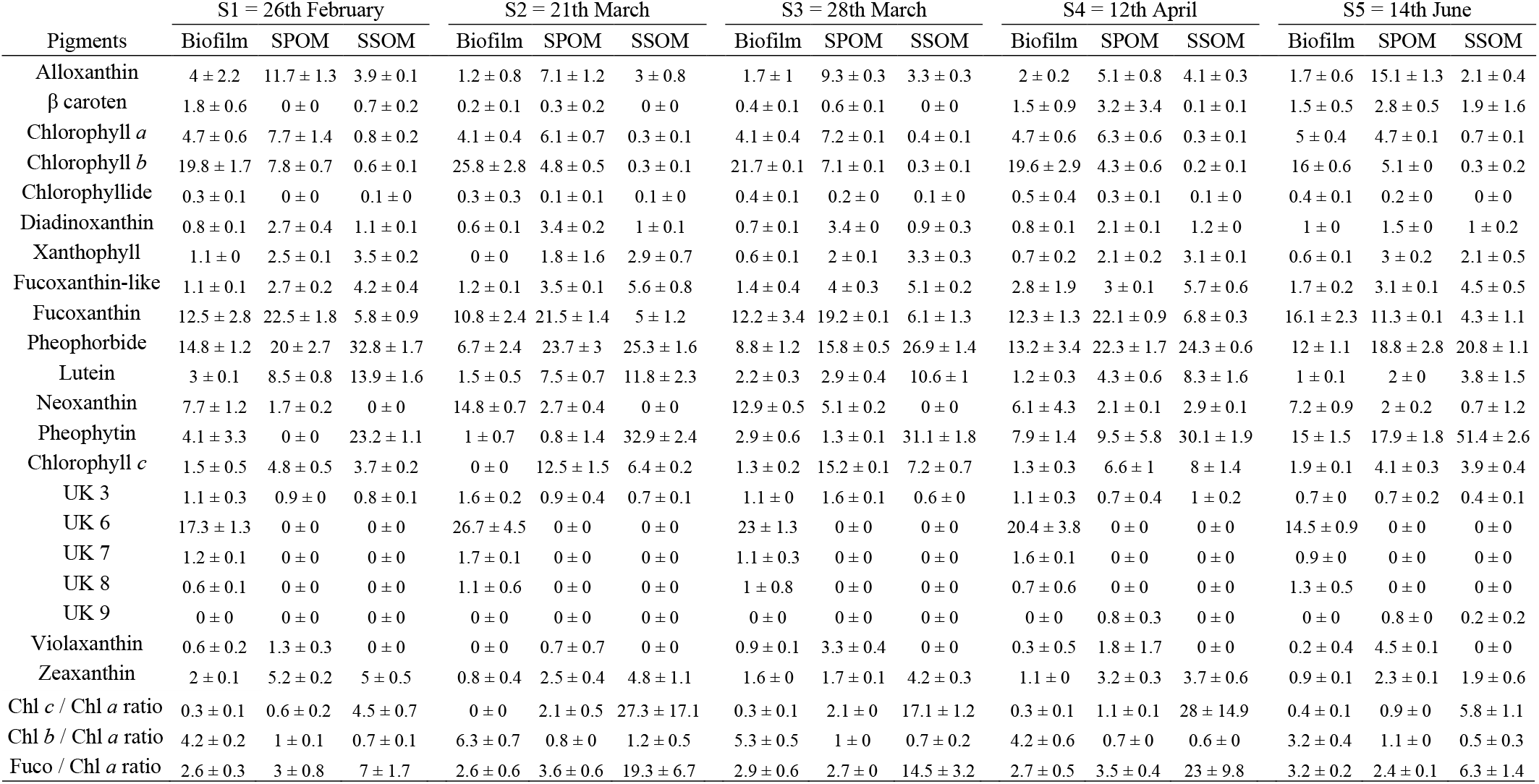
Pigment (%, mean ± SD, n ≥ 2) composition of organic matter sources (Biofilm, suspended particulate organic matter (SPOM), superficial sedimentary organic matter (SSOM)) over the sampling survey. UK: Unknown pigments.

| Pigments | S1 = 26th February |  |  | S2 = 21th March |  |  | S3 = 28th March |  |  | S4 = 12th April |  |  | S5 = 14th June |  |  |
| --- | --- | --- | --- | --- | --- | --- | --- | --- | --- | --- | --- | --- | --- | --- | --- |
|  | Biofilm | SPOM | SSOM | Biofilm | SPOM | SSOM | Biofilm | SPOM | SSOM | Biofilm | SPOM | SSOM | Biofilm | SPOM | SSOM |
| Alloxanthin | 4 ± 2.2 | 11.7 ± 1.3 | 3.9 ± 0.1 | 1.2 ± 0.8 | 7.1 ± 1.2 | 3 ± 0.8 | 1.7 ± 1 | 9.3 ± 0.3 | 3.3 ± 0.3 | 2 ± 0.2 | 5.1 ± 0.8 | 4.1 ± 0.3 | 1.7 ± 0.6 | 15.1 ± 1.3 | 2.1 ± 0.4 |
| $\beta$ caroten | 1.8 ± 0.6 | 0 ± 0 | 0.7 ± 0.2 | 0.2 ± 0.1 | 0.3 ± 0.2 | 0 ± 0 | 0.4 ± 0.1 | 0.6 ± 0.1 | 0 ± 0 | 1.5 ± 0.9 | 3.2 ± 3.4 | 0.1 ± 0.1 | 1.5 ± 0.5 | 2.8 ± 0.5 | 1.9 ± 1.6 |
| Chlorophyll *a* | 4.7 ± 0.6 | 7.7 ± 1.4 | 0.8 ± 0.2 | 4.1 ± 0.4 | 6.1 ± 0.7 | 0.3 ± 0.1 | 4.1 ± 0.4 | 7.2 ± 0.1 | 0.4 ± 0.1 | 4.7 ± 0.6 | 6.3 ± 0.6 | 0.3 ± 0.1 | 5 ± 0.4 | 4.7 ± 0.1 | 0.7 ± 0.1 |
| Chlorophyll *b* | 19.8 ± 1.7 | 7.8 ± 0.7 | 0.6 ± 0.1 | 25.8 ± 2.8 | 4.8 ± 0.5 | 0.3 ± 0.1 | 21.7 ± 0.1 | 7.1 ± 0.1 | 0.3 ± 0.1 | 19.6 ± 2.9 | 4.3 ± 0.6 | 0.2 ± 0.1 | 16 ± 0.6 | 5.1 ± 0 | 0.3 ± 0.2 |
| Chlorophyllide | 0.3 ± 0.1 | 0 ± 0 | 0.1 ± 0 | 0.3 ± 0.3 | 0.1 ± 0.1 | 0.1 ± 0 | 0.4 ± 0.1 | 0.2 ± 0 | 0.1 ± 0 | 0.5 ± 0.4 | 0.3 ± 0.1 | 0.1 ± 0 | 0.4 ± 0.1 | 0.2 ± 0 | 0 ± 0 |
| Diadinoxanthin | 0.8 ± 0.1 | 2.7 ± 0.4 | 1.1 ± 0.1 | 0.6 ± 0.1 | 3.4 ± 0.2 | 1 ± 0.1 | 0.7 ± 0.1 | 3.4 ± 0 | 0.9 ± 0.3 | 0.8 ± 0.1 | 2.1 ± 0.1 | 1.2 ± 0 | 1 ± 0 | 1.5 ± 0 | 1 ± 0.2 |
| Xanthophyll | 1.1 ± 0 | 2.5 ± 0.1 | 3.5 ± 0.2 | 0 ± 0 | 1.8 ± 1.6 | 2.9 ± 0.7 | 0.6 ± 0.1 | 2 ± 0.1 | 3.3 ± 0.3 | 0.7 ± 0.2 | 2.1 ± 0.2 | 3.1 ± 0.1 | 0.6 ± 0.1 | 3 ± 0.2 | 2.1 ± 0.5 |
| Fucoxanthin-like | 1.1 ± 0.1 | 2.7 ± 0.2 | 4.2 ± 0.4 | 1.2 ± 0.1 | 3.5 ± 0.1 | 5.6 ± 0.8 | 1.4 ± 0.4 | 4 ± 0.3 | 5.1 ± 0.2 | 2.8 ± 1.9 | 3 ± 0.1 | 5.7 ± 0.6 | 1.7 ± 0.2 | 3.1 ± 0.1 | 4.5 ± 0.5 |
| Fucoxanthin | 12.5 ± 2.8 | 22.5 ± 1.8 | 5.8 ± 0.9 | 10.8 ± 2.4 | 21.5 ± 1.4 | 5 ± 1.2 | 12.2 ± 3.4 | 19.2 ± 0.1 | 6.1 ± 1.3 | 12.3 ± 1.3 | 22.1 ± 0.9 | 6.8 ± 0.3 | 16.1 ± 2.3 | 11.3 ± 0.1 | 4.3 ± 1.1 |
| Pheophorbide | 14.8 ± 1.2 | 20 ± 2.7 | 32.8 ± 1.7 | 6.7 ± 2.4 | 23.7 ± 3 | 25.3 ± 1.6 | 8.8 ± 1.2 | 15.8 ± 0.5 | 26.9 ± 1.4 | 13.2 ± 3.4 | 22.3 ± 1.7 | 24.3 ± 0.6 | 12 ± 1.1 | 18.8 ± 2.8 | 20.8 ± 1.1 |
| Lutein | 3 ± 0.1 | 8.5 ± 0.8 | 13.9 ± 1.6 | 1.5 ± 0.5 | 7.5 ± 0.7 | 11.8 ± 2.3 | 2.2 ± 0.3 | 2.9 ± 0.4 | 10.6 ± 1 | 1.2 ± 0.3 | 4.3 ± 0.6 | 8.3 ± 1.6 | 1 ± 0.1 | 2 ± 0 | 3.8 ± 1.5 |
| Neoxanthin | 7.7 ± 1.2 | 1.7 ± 0.2 | 0 ± 0 | 14.8 ± 0.7 | 2.7 ± 0.4 | 0 ± 0 | 12.9 ± 0.5 | 5.1 ± 0.2 | 0 ± 0 | 6.1 ± 4.3 | 2.1 ± 0.1 | 2.9 ± 0.1 | 7.2 ± 0.9 | 2 ± 0.2 | 0.7 ± 1.2 |
| Pheophytin | 4.1 ± 3.3 | 0 ± 0 | 23.2 ± 1.1 | 1 ± 0.7 | 0.8 ± 1.4 | 32.9 ± 2.4 | 2.9 ± 0.6 | 1.3 ± 0.1 | 31.1 ± 1.8 | 7.9 ± 1.4 | 9.5 ± 5.8 | 30.1 ± 1.9 | 15 ± 1.5 | 17.9 ± 1.8 | 51.4 ± 2.6 |
| Chlorophyll *c* | 1.5 ± 0.5 | 4.8 ± 0.5 | 3.7 ± 0.2 | 0 ± 0 | 12.5 ± 1.5 | 6.4 ± 0.2 | 1.3 ± 0.2 | 15.2 ± 0.1 | 7.2 ± 0.7 | 1.3 ± 0.3 | 6.6 ± 1 | 8 ± 1.4 | 1.9 ± 0.1 | 4.1 ± 0.3 | 3.9 ± 0.4 |
| UK 3 | 1.1 ± 0.3 | 0.9 ± 0 | 0.8 ± 0.1 | 1.6 ± 0.2 | 0.9 ± 0.4 | 0.7 ± 0.1 | 1.1 ± 0 | 1.6 ± 0.1 | 0.6 ± 0 | 1.1 ± 0.3 | 0.7 ± 0.4 | 1 ± 0.2 | 0.7 ± 0 | 0.7 ± 0.2 | 0.4 ± 0.1 |
| UK 6 | 17.3 ± 1.3 | 0 ± 0 | 0 ± 0 | 26.7 ± 4.5 | 0 ± 0 | 0 ± 0 | 23 ± 1.3 | 0 ± 0 | 0 ± 0 | 20.4 ± 3.8 | 0 ± 0 | 0 ± 0 | 14.5 ± 0.9 | 0 ± 0 | 0 ± 0 |
| UK 7 | 1.2 ± 0.1 | 0 ± 0 | 0 ± 0 | 1.7 ± 0.1 | 0 ± 0 | 0 ± 0 | 1.1 ± 0.3 | 0 ± 0 | 0 ± 0 | 1.6 ± 0.1 | 0 ± 0 | 0 ± 0 | 0.9 ± 0 | 0 ± 0 | 0 ± 0 |
| UK 8 | 0.6 ± 0.1 | 0 ± 0 | 0 ± 0 | 1.1 ± 0.6 | 0 ± 0 | 0 ± 0 | 1 ± 0.8 | 0 ± 0 | 0 ± 0 | 0.7 ± 0.6 | 0 ± 0 | 0 ± 0 | 1.3 ± 0.5 | 0 ± 0 | 0 ± 0 |
| UK 9 | 0 ± 0 | 0 ± 0 | 0 ± 0 | 0 ± 0 | 0 ± 0 | 0 ± 0 | 0 ± 0 | 0 ± 0 | 0 ± 0 | 0 ± 0 | 0.8 ± 0.3 | 0 ± 0 | 0 ± 0 | 0.8 ± 0 | 0.2 ± 0.2 |
| Violaxanthin | 0.6 ± 0.2 | 1.3 ± 0.3 | 0 ± 0 | 0 ± 0 | 0.7 ± 0.7 | 0 ± 0 | 0.9 ± 0.1 | 3.3 ± 0.4 | 0 ± 0 | 0.3 ± 0.5 | 1.8 ± 1.7 | 0 ± 0 | 0.2 ± 0.4 | 4.5 ± 0.1 | 0 ± 0 |
| Zeaxanthin | 2 ± 0.1 | 5.2 ± 0.2 | 5 ± 0.5 | 0.8 ± 0.4 | 2.5 ± 0.4 | 4.8 ± 1.1 | 1.6 ± 0 | 1.7 ± 0.1 | 4.2 ± 0.3 | 1.1 ± 0 | 3.2 ± 0.3 | 3.7 ± 0.6 | 0.9 ± 0.1 | 2.3 ± 0.1 | 1.9 ± 0.6 |
| Chl *c* / Chl *a* ratio | 0.3 ± 0.1 | 0.6 ± 0.2 | 4.5 ± 0.7 | 0 ± 0 | 2.1 ± 0.5 | 27.3 ± 17.1 | 0.3 ± 0.1 | 2.1 ± 0 | 17.1 ± 1.2 | 0.3 ± 0.1 | 1.1 ± 0.1 | 28 ± 14.9 | 0.4 ± 0.1 | 0.9 ± 0 | 5.8 ± 1.1 |
| Chl *b* / Chl *a* ratio | 4.2 ± 0.2 | 1 ± 0.1 | 0.7 ± 0.1 | 6.3 ± 0.7 | 0.8 ± 0 | 1.2 ± 0.5 | 5.3 ± 0.5 | 1 ± 0 | 0.7 ± 0.2 | 4.2 ± 0.6 | 0.7 ± 0 | 0.6 ± 0 | 3.2 ± 0.4 | 1.1 ± 0 | 0.5 ± 0.3 |
| Fuco / Chl *a* ratio | 2.6 ± 0.3 | 3 ± 0.8 | 7 ± 1.7 | 2.6 ± 0.6 | 3.6 ± 0.6 | 19.3 ± 6.7 | 2.9 ± 0.6 | 2.7 ± 0 | 14.5 ± 3.2 | 2.7 ± 0.5 | 3.5 ± 0.4 | 23 ± 9.8 | 3.2 ± 0.2 | 2.4 ± 0.1 | 6.3 ± 1.4 |

**Table S2:**
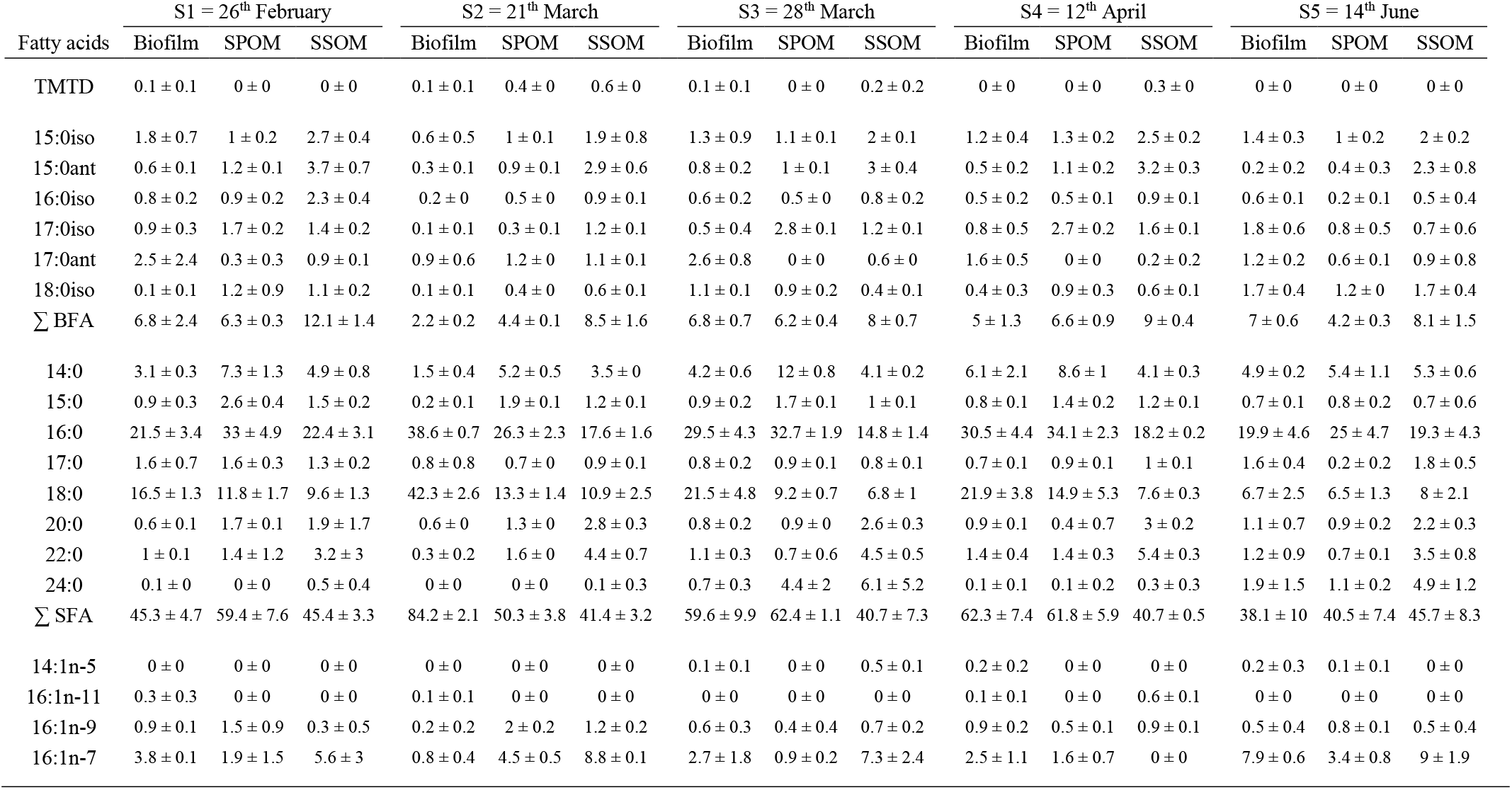
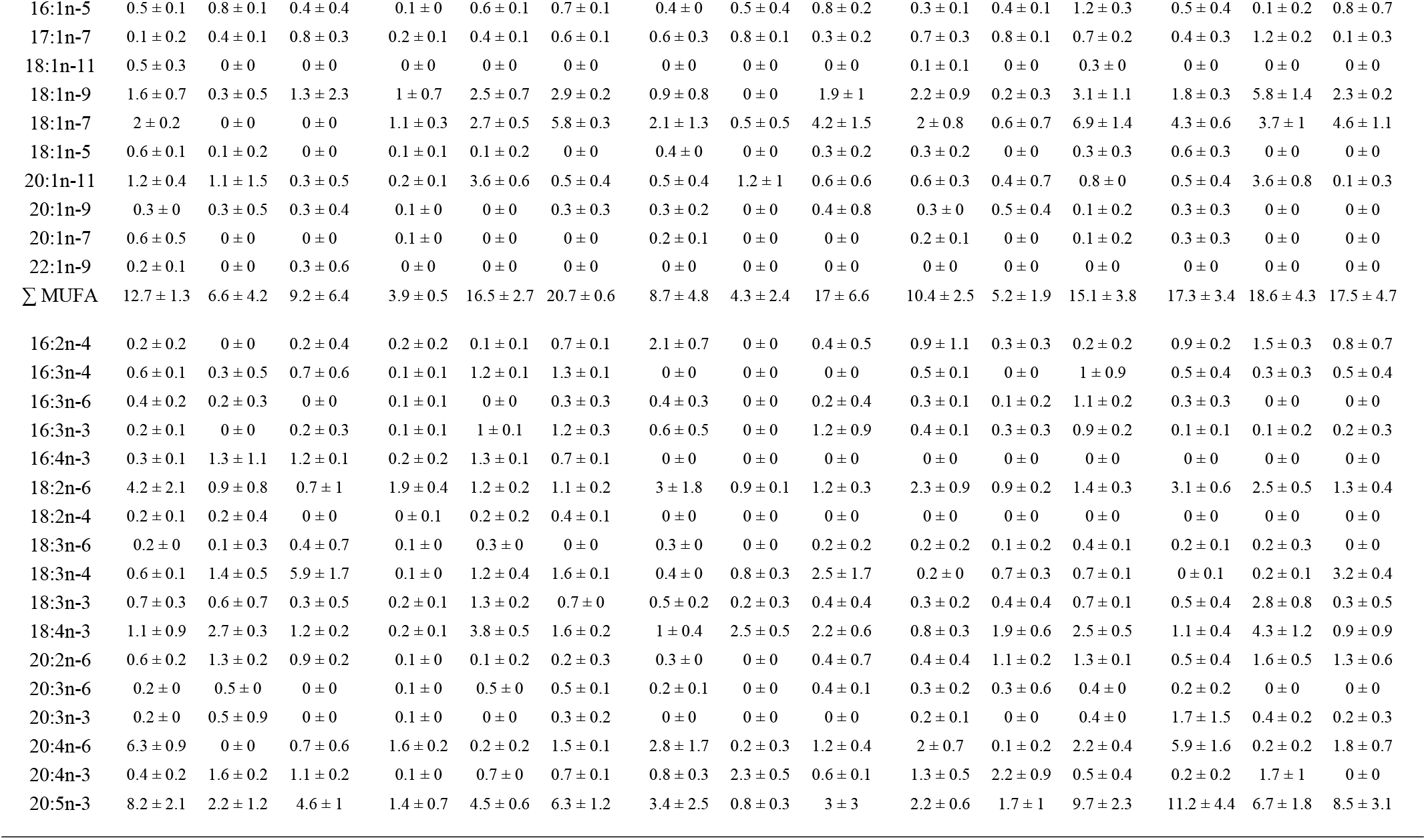
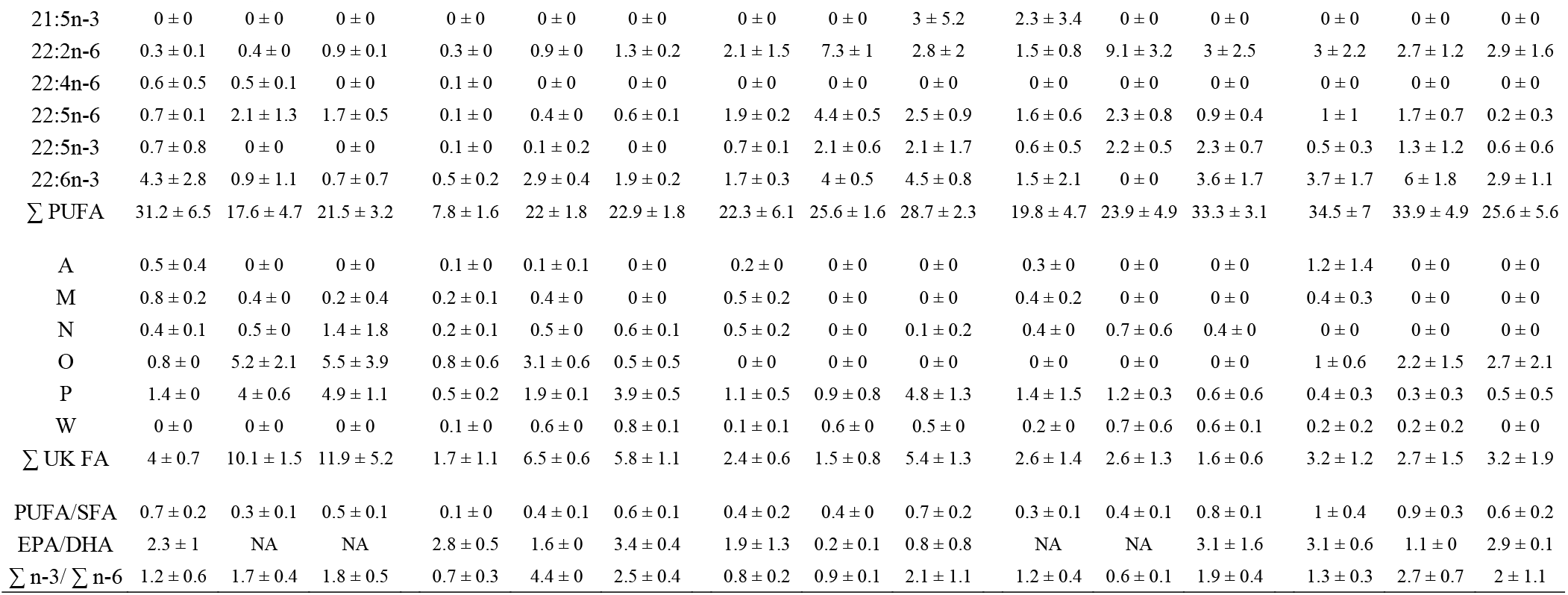
Fatty acid (FA) (%, mean ± SD, n = 3) composition of organic matter sources (Biofilm, suspended particulate organic matter (SPOM), superficial sedimentary organic matter (SSOM)) over the sampling survey. Only FA accounting for more than 0.5 % of total FA in at least one sample was shown. BFA: Branched FA; SFA: saturated FA; MUFA: monounsaturated FA; PUFA: polyunsaturated FA; UK FA: Unknown FA; EPA: 20:5n-3; DHA: 22:6n-3.

| Fatty acids | S1 = 26<sup>th</sup> February |  |  | S2 = 21<sup>th</sup> March |  |  | S3 = 28<sup>th</sup> March |  |  | S4 = 12<sup>th</sup> April |  |  | S5 = 14<sup>th</sup> June |  |  |
| --- | --- | --- | --- | --- | --- | --- | --- | --- | --- | --- | --- | --- | --- | --- | --- |
|  | Biofilm | SPOM | SSOM | Biofilm | SPOM | SSOM | Biofilm | SPOM | SSOM | Biofilm | SPOM | SSOM | Biofilm | SPOM | SSOM |
| TMTD | 0.1 ± 0.1 | 0 ± 0 | 0 ± 0 | 0.1 ± 0.1 | 0.4 ± 0 | 0.6 ± 0 | 0.1 ± 0.1 | 0 ± 0 | 0.2 ± 0.2 | 0 ± 0 | 0 ± 0 | 0.3 ± 0 | 0 ± 0 | 0 ± 0 | 0 ± 0 |
| 15:0iso | 1.8 ± 0.7 | 1 ± 0.2 | 2.7 ± 0.4 | 0.6 ± 0.5 | 1 ± 0.1 | 1.9 ± 0.8 | 1.3 ± 0.9 | 1.1 ± 0.1 | 2 ± 0.1 | 1.2 ± 0.4 | 1.3 ± 0.2 | 2.5 ± 0.2 | 1.4 ± 0.3 | 1 ± 0.2 | 2 ± 0.2 |
| 15:0ant | 0.6 ± 0.1 | 1.2 ± 0.1 | 3.7 ± 0.7 | 0.3 ± 0.1 | 0.9 ± 0.1 | 2.9 ± 0.6 | 0.8 ± 0.2 | 1 ± 0.1 | 3 ± 0.4 | 0.5 ± 0.2 | 1.1 ± 0.2 | 3.2 ± 0.3 | 0.2 ± 0.2 | 0.4 ± 0.3 | 2.3 ± 0.8 |
| 16:0iso | 0.8 ± 0.2 | 0.9 ± 0.2 | 2.3 ± 0.4 | 0.2 ± 0 | 0.5 ± 0 | 0.9 ± 0.1 | 0.6 ± 0.2 | 0.5 ± 0 | 0.8 ± 0.2 | 0.5 ± 0.2 | 0.5 ± 0.1 | 0.9 ± 0.1 | 0.6 ± 0.1 | 0.2 ± 0.1 | 0.5 ± 0.4 |
| 17:0iso | 0.9 ± 0.3 | 1.7 ± 0.2 | 1.4 ± 0.2 | 0.1 ± 0.1 | 0.3 ± 0.1 | 1.2 ± 0.1 | 0.5 ± 0.4 | 2.8 ± 0.1 | 1.2 ± 0.1 | 0.8 ± 0.5 | 2.7 ± 0.2 | 1.6 ± 0.1 | 1.8 ± 0.6 | 0.8 ± 0.5 | 0.7 ± 0.6 |
| 17:0ant | 2.5 ± 2.4 | 0.3 ± 0.3 | 0.9 ± 0.1 | 0.9 ± 0.6 | 1.2 ± 0 | 1.1 ± 0.1 | 2.6 ± 0.8 | 0 ± 0 | 0.6 ± 0 | 1.6 ± 0.5 | 0 ± 0 | 0.2 ± 0.2 | 1.2 ± 0.2 | 0.6 ± 0.1 | 0.9 ± 0.8 |
| 18:0iso | 0.1 ± 0.1 | 1.2 ± 0.9 | 1.1 ± 0.2 | 0.1 ± 0.1 | 0.4 ± 0 | 0.6 ± 0.1 | 1.1 ± 0.1 | 0.9 ± 0.2 | 0.4 ± 0.1 | 0.4 ± 0.3 | 0.9 ± 0.3 | 0.6 ± 0.1 | 1.7 ± 0.4 | 1.2 ± 0 | 1.7 ± 0.4 |
| $\Sigma$ BFA | 6.8 ± 2.4 | 6.3 ± 0.3 | 12.1 ± 1.4 | 2.2 ± 0.2 | 4.4 ± 0.1 | 8.5 ± 1.6 | 6.8 ± 0.7 | 6.2 ± 0.4 | 8 ± 0.7 | 5 ± 1.3 | 6.6 ± 0.9 | 9 ± 0.4 | 7 ± 0.6 | 4.2 ± 0.3 | 8.1 ± 1.5 |
| 14:0 | 3.1 ± 0.3 | 7.3 ± 1.3 | 4.9 ± 0.8 | 1.5 ± 0.4 | 5.2 ± 0.5 | 3.5 ± 0 | 4.2 ± 0.6 | 12 ± 0.8 | 4.1 ± 0.2 | 6.1 ± 2.1 | 8.6 ± 1 | 4.1 ± 0.3 | 4.9 ± 0.2 | 5.4 ± 1.1 | 5.3 ± 0.6 |
| 15:0 | 0.9 ± 0.3 | 2.6 ± 0.4 | 1.5 ± 0.2 | 0.2 ± 0.1 | 1.9 ± 0.1 | 1.2 ± 0.1 | 0.9 ± 0.2 | 1.7 ± 0.1 | 1 ± 0.1 | 0.8 ± 0.1 | 1.4 ± 0.2 | 1.2 ± 0.1 | 0.7 ± 0.1 | 0.8 ± 0.2 | 0.7 ± 0.6 |
| 16:0 | 21.5 ± 3.4 | 33 ± 4.9 | 22.4 ± 3.1 | 38.6 ± 0.7 | 26.3 ± 2.3 | 17.6 ± 1.6 | 29.5 ± 4.3 | 32.7 ± 1.9 | 14.8 ± 1.4 | 30.5 ± 4.4 | 34.1 ± 2.3 | 18.2 ± 0.2 | 19.9 ± 4.6 | 25 ± 4.7 | 19.3 ± 4.3 |
| 17:0 | 1.6 ± 0.7 | 1.6 ± 0.3 | 1.3 ± 0.2 | 0.8 ± 0.8 | 0.7 ± 0 | 0.9 ± 0.1 | 0.8 ± 0.2 | 0.9 ± 0.1 | 0.8 ± 0.1 | 0.7 ± 0.1 | 0.9 ± 0.1 | 1 ± 0.1 | 1.6 ± 0.4 | 0.2 ± 0.2 | 1.8 ± 0.5 |
| 18:0 | 16.5 ± 1.3 | 11.8 ± 1.7 | 9.6 ± 1.3 | 42.3 ± 2.6 | 13.3 ± 1.4 | 10.9 ± 2.5 | 21.5 ± 4.8 | 9.2 ± 0.7 | 6.8 ± 1 | 21.9 ± 3.8 | 14.9 ± 5.3 | 7.6 ± 0.3 | 6.7 ± 2.5 | 6.5 ± 1.3 | 8 ± 2.1 |
| 20:0 | 0.6 ± 0.1 | 1.7 ± 0.1 | 1.9 ± 1.7 | 0.6 ± 0 | 1.3 ± 0 | 2.8 ± 0.3 | 0.8 ± 0.2 | 0.9 ± 0 | 2.6 ± 0.3 | 0.9 ± 0.1 | 0.4 ± 0.7 | 3 ± 0.2 | 1.1 ± 0.7 | 0.9 ± 0.2 | 2.2 ± 0.3 |
| 22:0 | 1 ± 0.1 | 1.4 ± 1.2 | 3.2 ± 3 | 0.3 ± 0.2 | 1.6 ± 0 | 4.4 ± 0.7 | 1.1 ± 0.3 | 0.7 ± 0.6 | 4.5 ± 0.5 | 1.4 ± 0.4 | 1.4 ± 0.3 | 5.4 ± 0.3 | 1.2 ± 0.9 | 0.7 ± 0.1 | 3.5 ± 0.8 |
| 24:0 | 0.1 ± 0 | 0 ± 0 | 0.5 ± 0.4 | 0 ± 0 | 0 ± 0 | 0.1 ± 0.3 | 0.7 ± 0.3 | 4.4 ± 2 | 6.1 ± 5.2 | 0.1 ± 0.1 | 0.1 ± 0.2 | 0.3 ± 0.3 | 1.9 ± 1.5 | 1.1 ± 0.2 | 4.9 ± 1.2 |
| $\Sigma$ SFA | 45.3 ± 4.7 | 59.4 ± 7.6 | 45.4 ± 3.3 | 84.2 ± 2.1 | 50.3 ± 3.8 | 41.4 ± 3.2 | 59.6 ± 9.9 | 62.4 ± 1.1 | 40.7 ± 7.3 | 62.3 ± 7.4 | 61.8 ± 5.9 | 40.7 ± 0.5 | 38.1 ± 10 | 40.5 ± 7.4 | 45.7 ± 8.3 |
| 14:1n-5 | 0 ± 0 | 0 ± 0 | 0 ± 0 | 0 ± 0 | 0 ± 0 | 0 ± 0 | 0.1 ± 0.1 | 0 ± 0 | 0.5 ± 0.1 | 0.2 ± 0.2 | 0 ± 0 | 0 ± 0 | 0.2 ± 0.3 | 0.1 ± 0.1 | 0 ± 0 |
| 16:1n-11 | 0.3 ± 0.3 | 0 ± 0 | 0 ± 0 | 0.1 ± 0.1 | 0 ± 0 | 0 ± 0 | 0 ± 0 | 0 ± 0 | 0 ± 0 | 0.1 ± 0.1 | 0 ± 0 | 0.6 ± 0.1 | 0 ± 0 | 0 ± 0 | 0 ± 0 |
| 16:1n-9 | 0.9 ± 0.1 | 1.5 ± 0.9 | 0.3 ± 0.5 | 0.2 ± 0.2 | 2 ± 0.2 | 1.2 ± 0.2 | 0.6 ± 0.3 | 0.4 ± 0.4 | 0.7 ± 0.2 | 0.9 ± 0.2 | 0.5 ± 0.1 | 0.9 ± 0.1 | 0.5 ± 0.4 | 0.8 ± 0.1 | 0.5 ± 0.4 |
| 16:1n-7 | 3.8 ± 0.1 | 1.9 ± 1.5 | 5.6 ± 3 | 0.8 ± 0.4 | 4.5 ± 0.5 | 8.8 ± 0.1 | 2.7 ± 1.8 | 0.9 ± 0.2 | 7.3 ± 2.4 | 2.5 ± 1.1 | 1.6 ± 0.7 | 0 ± 0 | 7.9 ± 0.6 | 3.4 ± 0.8 | 9 ± 1.9 |
| 16:1n-5 | 0.5 ± 0.1 | 0.8 ± 0.1 | 0.4 ± 0.4 | 0.1 ± 0 | 0.6 ± 0.1 | 0.7 ± 0.1 | 0.4 ± 0 | 0.5 ± 0.4 | 0.8 ± 0.2 | 0.3 ± 0.1 | 0.4 ± 0.1 | 1.2 ± 0.3 | 0.5 ± 0.4 | 0.1 ± 0.2 | 0.8 ± 0.7 |
| 17:1n-7 | 0.1 ± 0.2 | 0.4 ± 0.1 | 0.8 ± 0.3 | 0.2 ± 0.1 | 0.4 ± 0.1 | 0.6 ± 0.1 | 0.6 ± 0.3 | 0.8 ± 0.1 | 0.3 ± 0.2 | 0.7 ± 0.3 | 0.8 ± 0.1 | 0.7 ± 0.2 | 0.4 ± 0.3 | 1.2 ± 0.2 | 0.1 ± 0.3 |
| 18:1n-11 | 0.5 ± 0.3 | 0 ± 0 | 0 ± 0 | 0 ± 0 | 0 ± 0 | 0 ± 0 | 0 ± 0 | 0 ± 0 | 0 ± 0 | 0.1 ± 0.1 | 0 ± 0 | 0.3 ± 0 | 0 ± 0 | 0 ± 0 | 0 ± 0 |
| 18:1n-9 | 1.6 ± 0.7 | 0.3 ± 0.5 | 1.3 ± 2.3 | 1 ± 0.7 | 2.5 ± 0.7 | 2.9 ± 0.2 | 0.9 ± 0.8 | 0 ± 0 | 1.9 ± 1 | 2.2 ± 0.9 | 0.2 ± 0.3 | 3.1 ± 1.1 | 1.8 ± 0.3 | 5.8 ± 1.4 | 2.3 ± 0.2 |
| 18:1n-7 | 2 ± 0.2 | 0 ± 0 | 0 ± 0 | 1.1 ± 0.3 | 2.7 ± 0.5 | 5.8 ± 0.3 | 2.1 ± 1.3 | 0.5 ± 0.5 | 4.2 ± 1.5 | 2 ± 0.8 | 0.6 ± 0.7 | 6.9 ± 1.4 | 4.3 ± 0.6 | 3.7 ± 1 | 4.6 ± 1.1 |
| 18:1n-5 | 0.6 ± 0.1 | 0.1 ± 0.2 | 0 ± 0 | 0.1 ± 0.1 | 0.1 ± 0.2 | 0 ± 0 | 0.4 ± 0 | 0 ± 0 | 0.3 ± 0.2 | 0.3 ± 0.2 | 0 ± 0 | 0.3 ± 0.3 | 0.6 ± 0.3 | 0 ± 0 | 0 ± 0 |
| 20:1n-11 | 1.2 ± 0.4 | 1.1 ± 1.5 | 0.3 ± 0.5 | 0.2 ± 0.1 | 3.6 ± 0.6 | 0.5 ± 0.4 | 0.5 ± 0.4 | 1.2 ± 1 | 0.6 ± 0.6 | 0.6 ± 0.3 | 0.4 ± 0.7 | 0.8 ± 0 | 0.5 ± 0.4 | 3.6 ± 0.8 | 0.1 ± 0.3 |
| 20:1n-9 | 0.3 ± 0 | 0.3 ± 0.5 | 0.3 ± 0.4 | 0.1 ± 0 | 0 ± 0 | 0.3 ± 0.3 | 0.3 ± 0.2 | 0 ± 0 | 0.4 ± 0.8 | 0.3 ± 0 | 0.5 ± 0.4 | 0.1 ± 0.2 | 0.3 ± 0.3 | 0 ± 0 | 0 ± 0 |
| 20:1n-7 | 0.6 ± 0.5 | 0 ± 0 | 0 ± 0 | 0.1 ± 0 | 0 ± 0 | 0 ± 0 | 0.2 ± 0.1 | 0 ± 0 | 0 ± 0 | 0.2 ± 0.1 | 0 ± 0 | 0.1 ± 0.2 | 0.3 ± 0.3 | 0 ± 0 | 0 ± 0 |
| 22:1n-9 | 0.2 ± 0.1 | 0 ± 0 | 0.3 ± 0.6 | 0 ± 0 | 0 ± 0 | 0 ± 0 | 0 ± 0 | 0 ± 0 | 0 ± 0 | 0 ± 0 | 0 ± 0 | 0 ± 0 | 0 ± 0 | 0 ± 0 | 0 ± 0 |
| Σ MUFA | 12.7 ± 1.3 | 6.6 ± 4.2 | 9.2 ± 6.4 | 3.9 ± 0.5 | 16.5 ± 2.7 | 20.7 ± 0.6 | 8.7 ± 4.8 | 4.3 ± 2.4 | 17 ± 6.6 | 10.4 ± 2.5 | 5.2 ± 1.9 | 15.1 ± 3.8 | 17.3 ± 3.4 | 18.6 ± 4.3 | 17.5 ± 4.7 |
| 16:2n-4 | 0.2 ± 0.2 | 0 ± 0 | 0.2 ± 0.4 | 0.2 ± 0.2 | 0.1 ± 0.1 | 0.7 ± 0.1 | 2.1 ± 0.7 | 0 ± 0 | 0.4 ± 0.5 | 0.9 ± 1.1 | 0.3 ± 0.3 | 0.2 ± 0.2 | 0.9 ± 0.2 | 1.5 ± 0.3 | 0.8 ± 0.7 |
| 16:3n-4 | 0.6 ± 0.1 | 0.3 ± 0.5 | 0.7 ± 0.6 | 0.1 ± 0.1 | 1.2 ± 0.1 | 1.3 ± 0.1 | 0 ± 0 | 0 ± 0 | 0 ± 0 | 0.5 ± 0.1 | 0 ± 0 | 1 ± 0.9 | 0.5 ± 0.4 | 0.3 ± 0.3 | 0.5 ± 0.4 |
| 16:3n-6 | 0.4 ± 0.2 | 0.2 ± 0.3 | 0 ± 0 | 0.1 ± 0.1 | 0 ± 0 | 0.3 ± 0.3 | 0.4 ± 0.3 | 0 ± 0 | 0.2 ± 0.4 | 0.3 ± 0.1 | 0.1 ± 0.2 | 1.1 ± 0.2 | 0.3 ± 0.3 | 0 ± 0 | 0 ± 0 |
| 16:3n-3 | 0.2 ± 0.1 | 0 ± 0 | 0.2 ± 0.3 | 0.1 ± 0.1 | 1 ± 0.1 | 1.2 ± 0.3 | 0.6 ± 0.5 | 0 ± 0 | 1.2 ± 0.9 | 0.4 ± 0.1 | 0.3 ± 0.3 | 0.9 ± 0.2 | 0.1 ± 0.1 | 0.1 ± 0.2 | 0.2 ± 0.3 |
| 16:4n-3 | 0.3 ± 0.1 | 1.3 ± 1.1 | 1.2 ± 0.1 | 0.2 ± 0.2 | 1.3 ± 0.1 | 0.7 ± 0.1 | 0 ± 0 | 0 ± 0 | 0 ± 0 | 0 ± 0 | 0 ± 0 | 0 ± 0 | 0 ± 0 | 0 ± 0 | 0 ± 0 |
| 18:2n-6 | 4.2 ± 2.1 | 0.9 ± 0.8 | 0.7 ± 1 | 1.9 ± 0.4 | 1.2 ± 0.2 | 1.1 ± 0.2 | 3 ± 1.8 | 0.9 ± 0.1 | 1.2 ± 0.3 | 2.3 ± 0.9 | 0.9 ± 0.2 | 1.4 ± 0.3 | 3.1 ± 0.6 | 2.5 ± 0.5 | 1.3 ± 0.4 |
| 18:2n-4 | 0.2 ± 0.1 | 0.2 ± 0.4 | 0 ± 0 | 0 ± 0.1 | 0.2 ± 0.2 | 0.4 ± 0.1 | 0 ± 0 | 0 ± 0 | 0 ± 0 | 0 ± 0 | 0 ± 0 | 0 ± 0 | 0 ± 0 | 0 ± 0 | 0 ± 0 |
| 18:3n-6 | 0.2 ± 0 | 0.1 ± 0.3 | 0.4 ± 0.7 | 0.1 ± 0 | 0.3 ± 0 | 0 ± 0 | 0.3 ± 0 | 0 ± 0 | 0.2 ± 0.2 | 0.2 ± 0.2 | 0.1 ± 0.2 | 0.4 ± 0.1 | 0.2 ± 0.1 | 0.2 ± 0.3 | 0 ± 0 |
| 18:3n-4 | 0.6 ± 0.1 | 1.4 ± 0.5 | 5.9 ± 1.7 | 0.1 ± 0 | 1.2 ± 0.4 | 1.6 ± 0.1 | 0.4 ± 0 | 0.8 ± 0.3 | 2.5 ± 1.7 | 0.2 ± 0 | 0.7 ± 0.3 | 0.7 ± 0.1 | 0 ± 0.1 | 0.2 ± 0.1 | 3.2 ± 0.4 |
| 18:3n-3 | 0.7 ± 0.3 | 0.6 ± 0.7 | 0.3 ± 0.5 | 0.2 ± 0.1 | 1.3 ± 0.2 | 0.7 ± 0 | 0.5 ± 0.2 | 0.2 ± 0.3 | 0.4 ± 0.4 | 0.3 ± 0.2 | 0.4 ± 0.4 | 0.7 ± 0.1 | 0.5 ± 0.4 | 2.8 ± 0.8 | 0.3 ± 0.5 |
| 18:4n-3 | 1.1 ± 0.9 | 2.7 ± 0.3 | 1.2 ± 0.2 | 0.2 ± 0.1 | 3.8 ± 0.5 | 1.6 ± 0.2 | 1 ± 0.4 | 2.5 ± 0.5 | 2.2 ± 0.6 | 0.8 ± 0.3 | 1.9 ± 0.6 | 2.5 ± 0.5 | 1.1 ± 0.4 | 4.3 ± 1.2 | 0.9 ± 0.9 |
| 20:2n-6 | 0.6 ± 0.2 | 1.3 ± 0.2 | 0.9 ± 0.2 | 0.1 ± 0 | 0.1 ± 0.2 | 0.2 ± 0.3 | 0.3 ± 0 | 0 ± 0 | 0.4 ± 0.7 | 0.4 ± 0.4 | 1.1 ± 0.2 | 1.3 ± 0.1 | 0.5 ± 0.4 | 1.6 ± 0.5 | 1.3 ± 0.6 |
| 20:3n-6 | 0.2 ± 0 | 0.5 ± 0 | 0 ± 0 | 0.1 ± 0 | 0.5 ± 0 | 0.5 ± 0.1 | 0.2 ± 0.1 | 0 ± 0 | 0.4 ± 0.1 | 0.3 ± 0.2 | 0.3 ± 0.6 | 0.4 ± 0 | 0.2 ± 0.2 | 0 ± 0 | 0 ± 0 |
| 20:3n-3 | 0.2 ± 0 | 0.5 ± 0.9 | 0 ± 0 | 0.1 ± 0 | 0 ± 0 | 0.3 ± 0.2 | 0 ± 0 | 0 ± 0 | 0 ± 0 | 0.2 ± 0.1 | 0 ± 0 | 0.4 ± 0 | 1.7 ± 1.5 | 0.4 ± 0.2 | 0.2 ± 0.3 |
| 20:4n-6 | 6.3 ± 0.9 | 0 ± 0 | 0.7 ± 0.6 | 1.6 ± 0.2 | 0.2 ± 0.2 | 1.5 ± 0.1 | 2.8 ± 1.7 | 0.2 ± 0.3 | 1.2 ± 0.4 | 2 ± 0.7 | 0.1 ± 0.2 | 2.2 ± 0.4 | 5.9 ± 1.6 | 0.2 ± 0.2 | 1.8 ± 0.7 |
| 20:4n-3 | 0.4 ± 0.2 | 1.6 ± 0.2 | 1.1 ± 0.2 | 0.1 ± 0 | 0.7 ± 0 | 0.7 ± 0.1 | 0.8 ± 0.3 | 2.3 ± 0.5 | 0.6 ± 0.1 | 1.3 ± 0.5 | 2.2 ± 0.9 | 0.5 ± 0.4 | 0.2 ± 0.2 | 1.7 ± 1 | 0 ± 0 |
| 20:5n-3 | 8.2 ± 2.1 | 2.2 ± 1.2 | 4.6 ± 1 | 1.4 ± 0.7 | 4.5 ± 0.6 | 6.3 ± 1.2 | 3.4 ± 2.5 | 0.8 ± 0.3 | 3 ± 3 | 2.2 ± 0.6 | 1.7 ± 1 | 9.7 ± 2.3 | 11.2 ± 4.4 | 6.7 ± 1.8 | 8.5 ± 3.1 |
| 21:5n-3 | 0 ± 0 | 0 ± 0 | 0 ± 0 | 0 ± 0 | 0 ± 0 | 0 ± 0 | 0 ± 0 | 0 ± 0 | 3 ± 5.2 | 2.3 ± 3.4 | 0 ± 0 | 0 ± 0 | 0 ± 0 | 0 ± 0 | 0 ± 0 |
| 22:2n-6 | 0.3 ± 0.1 | 0.4 ± 0 | 0.9 ± 0.1 | 0.3 ± 0 | 0.9 ± 0 | 1.3 ± 0.2 | 2.1 ± 1.5 | 7.3 ± 1 | 2.8 ± 2 | 1.5 ± 0.8 | 9.1 ± 3.2 | 3 ± 2.5 | 3 ± 2.2 | 2.7 ± 1.2 | 2.9 ± 1.6 |
| 22:4n-6 | 0.6 ± 0.5 | 0.5 ± 0.1 | 0 ± 0 | 0.1 ± 0 | 0 ± 0 | 0 ± 0 | 0 ± 0 | 0 ± 0 | 0 ± 0 | 0 ± 0 | 0 ± 0 | 0 ± 0 | 0 ± 0 | 0 ± 0 | 0 ± 0 |
| 22:5n-6 | 0.7 ± 0.1 | 2.1 ± 1.3 | 1.7 ± 0.5 | 0.1 ± 0 | 0.4 ± 0 | 0.6 ± 0.1 | 1.9 ± 0.2 | 4.4 ± 0.5 | 2.5 ± 0.9 | 1.6 ± 0.6 | 2.3 ± 0.8 | 0.9 ± 0.4 | 1 ± 1 | 1.7 ± 0.7 | 0.2 ± 0.3 |
| 22:5n-3 | 0.7 ± 0.8 | 0 ± 0 | 0 ± 0 | 0.1 ± 0 | 0.1 ± 0.2 | 0 ± 0 | 0.7 ± 0.1 | 2.1 ± 0.6 | 2.1 ± 1.7 | 0.6 ± 0.5 | 2.2 ± 0.5 | 2.3 ± 0.7 | 0.5 ± 0.3 | 1.3 ± 1.2 | 0.6 ± 0.6 |
| 22:6n-3 | 4.3 ± 2.8 | 0.9 ± 1.1 | 0.7 ± 0.7 | 0.5 ± 0.2 | 2.9 ± 0.4 | 1.9 ± 0.2 | 1.7 ± 0.3 | 4 ± 0.5 | 4.5 ± 0.8 | 1.5 ± 2.1 | 0 ± 0 | 3.6 ± 1.7 | 3.7 ± 1.7 | 6 ± 1.8 | 2.9 ± 1.1 |
| Σ PUFA | 31.2 ± 6.5 | 17.6 ± 4.7 | 21.5 ± 3.2 | 7.8 ± 1.6 | 22 ± 1.8 | 22.9 ± 1.8 | 22.3 ± 6.1 | 25.6 ± 1.6 | 28.7 ± 2.3 | 19.8 ± 4.7 | 23.9 ± 4.9 | 33.3 ± 3.1 | 34.5 ± 7 | 33.9 ± 4.9 | 25.6 ± 5.6 |
| A | 0.5 ± 0.4 | 0 ± 0 | 0 ± 0 | 0.1 ± 0 | 0.1 ± 0.1 | 0 ± 0 | 0.2 ± 0 | 0 ± 0 | 0 ± 0 | 0.3 ± 0 | 0 ± 0 | 0 ± 0 | 1.2 ± 1.4 | 0 ± 0 | 0 ± 0 |
| M | 0.8 ± 0.2 | 0.4 ± 0 | 0.2 ± 0.4 | 0.2 ± 0.1 | 0.4 ± 0 | 0 ± 0 | 0.5 ± 0.2 | 0 ± 0 | 0 ± 0 | 0.4 ± 0.2 | 0 ± 0 | 0 ± 0 | 0.4 ± 0.3 | 0 ± 0 | 0 ± 0 |
| N | 0.4 ± 0.1 | 0.5 ± 0 | 1.4 ± 1.8 | 0.2 ± 0.1 | 0.5 ± 0 | 0.6 ± 0.1 | 0.5 ± 0.2 | 0 ± 0 | 0.1 ± 0.2 | 0.4 ± 0 | 0.7 ± 0.6 | 0.4 ± 0 | 0 ± 0 | 0 ± 0 | 0 ± 0 |
| O | 0.8 ± 0 | 5.2 ± 2.1 | 5.5 ± 3.9 | 0.8 ± 0.6 | 3.1 ± 0.6 | 0.5 ± 0.5 | 0 ± 0 | 0 ± 0 | 0 ± 0 | 0 ± 0 | 0 ± 0 | 0 ± 0 | 1 ± 0.6 | 2.2 ± 1.5 | 2.7 ± 2.1 |
| P | 1.4 ± 0 | 4 ± 0.6 | 4.9 ± 1.1 | 0.5 ± 0.2 | 1.9 ± 0.1 | 3.9 ± 0.5 | 1.1 ± 0.5 | 0.9 ± 0.8 | 4.8 ± 1.3 | 1.4 ± 1.5 | 1.2 ± 0.3 | 0.6 ± 0.6 | 0.4 ± 0.3 | 0.3 ± 0.3 | 0.5 ± 0.5 |
| W | 0 ± 0 | 0 ± 0 | 0 ± 0 | 0.1 ± 0 | 0.6 ± 0 | 0.8 ± 0.1 | 0.1 ± 0.1 | 0.6 ± 0 | 0.5 ± 0 | 0.2 ± 0 | 0.7 ± 0.6 | 0.6 ± 0.1 | 0.2 ± 0.2 | 0.2 ± 0.2 | 0 ± 0 |
| Σ UK FA | 4 ± 0.7 | 10.1 ± 1.5 | 11.9 ± 5.2 | 1.7 ± 1.1 | 6.5 ± 0.6 | 5.8 ± 1.1 | 2.4 ± 0.6 | 1.5 ± 0.8 | 5.4 ± 1.3 | 2.6 ± 1.4 | 2.6 ± 1.3 | 1.6 ± 0.6 | 3.2 ± 1.2 | 2.7 ± 1.5 | 3.2 ± 1.9 |
| PUFA/SFA | 0.7 ± 0.2 | 0.3 ± 0.1 | 0.5 ± 0.1 | 0.1 ± 0 | 0.4 ± 0.1 | 0.6 ± 0.1 | 0.4 ± 0.2 | 0.4 ± 0 | 0.7 ± 0.2 | 0.3 ± 0.1 | 0.4 ± 0.1 | 0.8 ± 0.1 | 1 ± 0.4 | 0.9 ± 0.3 | 0.6 ± 0.2 |
| EPA/DHA | 2.3 ± 1 | NA | NA | 2.8 ± 0.5 | 1.6 ± 0 | 3.4 ± 0.4 | 1.9 ± 1.3 | 0.2 ± 0.1 | 0.8 ± 0.8 | NA | NA | 3.1 ± 1.6 | 3.1 ± 0.6 | 1.1 ± 0 | 2.9 ± 0.1 |
| Σ n-3/ Σ n-6 | 1.2 ± 0.6 | 1.7 ± 0.4 | 1.8 ± 0.5 | 0.7 ± 0.3 | 4.4 ± 0 | 2.5 ± 0.4 | 0.8 ± 0.2 | 0.9 ± 0.1 | 2.1 ± 1.1 | 1.2 ± 0.4 | 0.6 ± 0.1 | 1.9 ± 0.4 | 1.3 ± 0.3 | 2.7 ± 0.7 | 2 ± 1.1 |

**Table S3:** Stable isotope (mean ± SD, n = 3) composition of organic matter sources (Biofilm, suspended particulate organic matter (SPOM), superficial sedimentary organic matter (SSOM)) over the sampling survey.

|  | S1 = 26<sup>th</sup> February |  |  | S2 = 21<sup>th</sup> March |  |  | S3 = 28<sup>th</sup> March |  |  | S4 = 12<sup>th</sup> April |  |  | S5 = 14<sup>th</sup> June |  |  |
| --- | --- | --- | --- | --- | --- | --- | --- | --- | --- | --- | --- | --- | --- | --- | --- |
|  | Biofilm | SPOM | SSOM | Biofilm | SPOM | SSOM | Biofilm | SPOM | SSOM | Biofilm | SPOM | SSOM | Biofilm | SPOM | SSOM |
| $\delta^{13}\text{C}$ | -22.6 ± 0.1 | -25.2 ± 0.2 | -23 ± 0.2 | -23.4 ± 0.5 | -25.3 ± 0.1 | -22.7 ± 0.3 | -22.6 ± 0.4 | -25.3 ± 0.1 | -22.6 ± 0.1 | -23.4 ± 1 | -24.6 ± 0.4 | -22.5 ± 0.1 | -21.8 ± 0.2 | -25.1 ± 0.3 | -22.5 ± 0.1 |
| $\delta^{15}\text{N}$ | 9.4 ± 0.6 | 7.3 ± 0 | 7 ± 0.7 | 10.3 ± 0.5 | 6.7 ± 0.3 | 7.1 ± 0.1 | 9.2 ± 0.7 | 6.6 ± 0.1 | 7 ± 0.1 | 9.2 ± 0.7 | 6.9 ± 0.9 | 7.1 ± 0.1 | 9.4 ± 0.7 | 6.4 ± 0.5 | 6.9 ± 0.2 |
| C/N | 3.9 ± 0.3 | 6.6 ± 1.1 | 5.1 ± 1.3 | 5.8 ± 3 | 5.9 ± 0.6 | 5.3 ± 0.6 | 5.3 ± 0.4 | 6 ± 0.5 | 5.5 ± 0.6 | 5 ± 0.5 | 5.6 ± 0.9 | 5.4 ± 0.5 | 3.6 ± 0.6 | 3.9 ± 0.4 | 5.9 ± 0.8 |

**Table S4:**
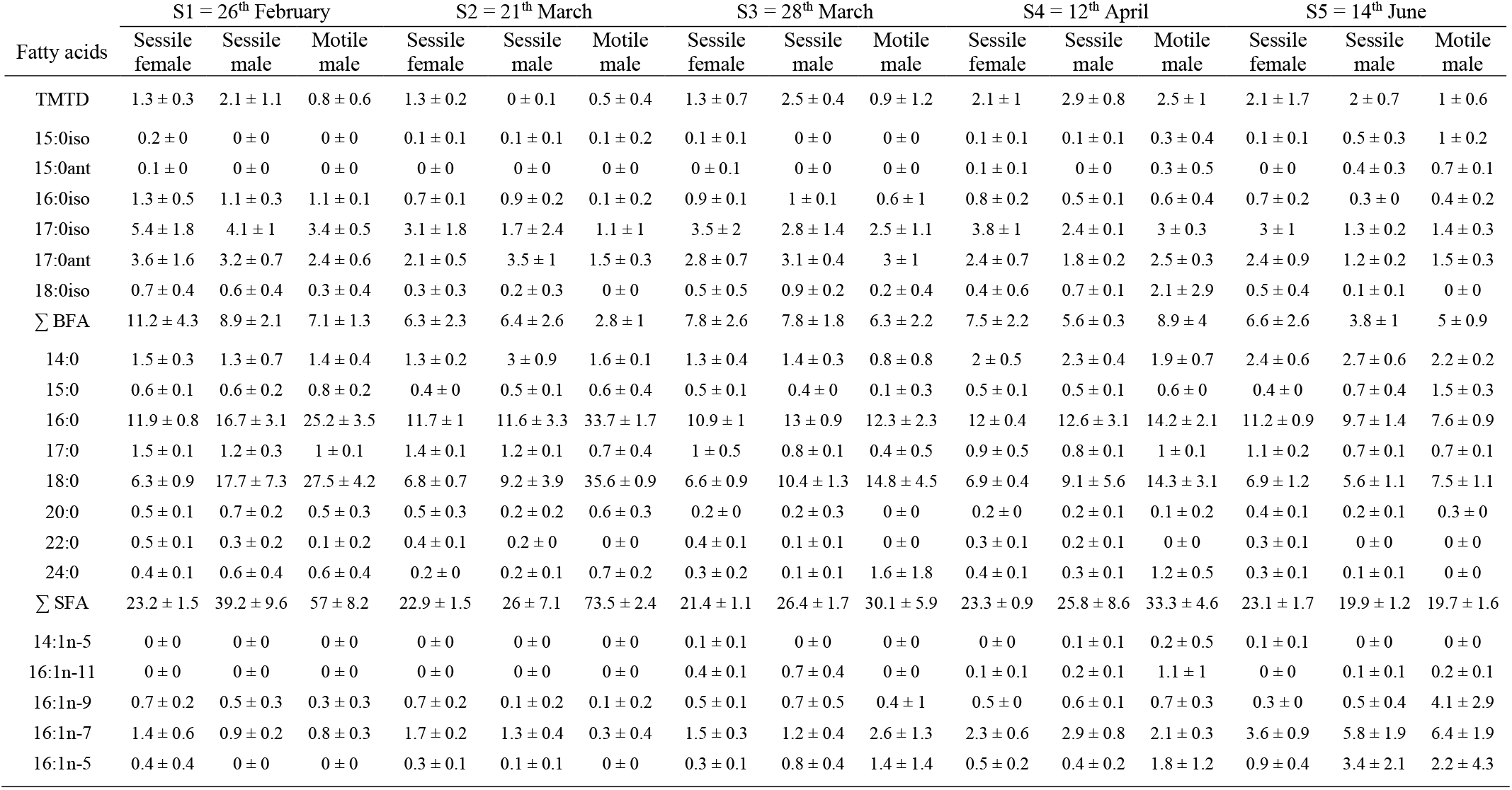
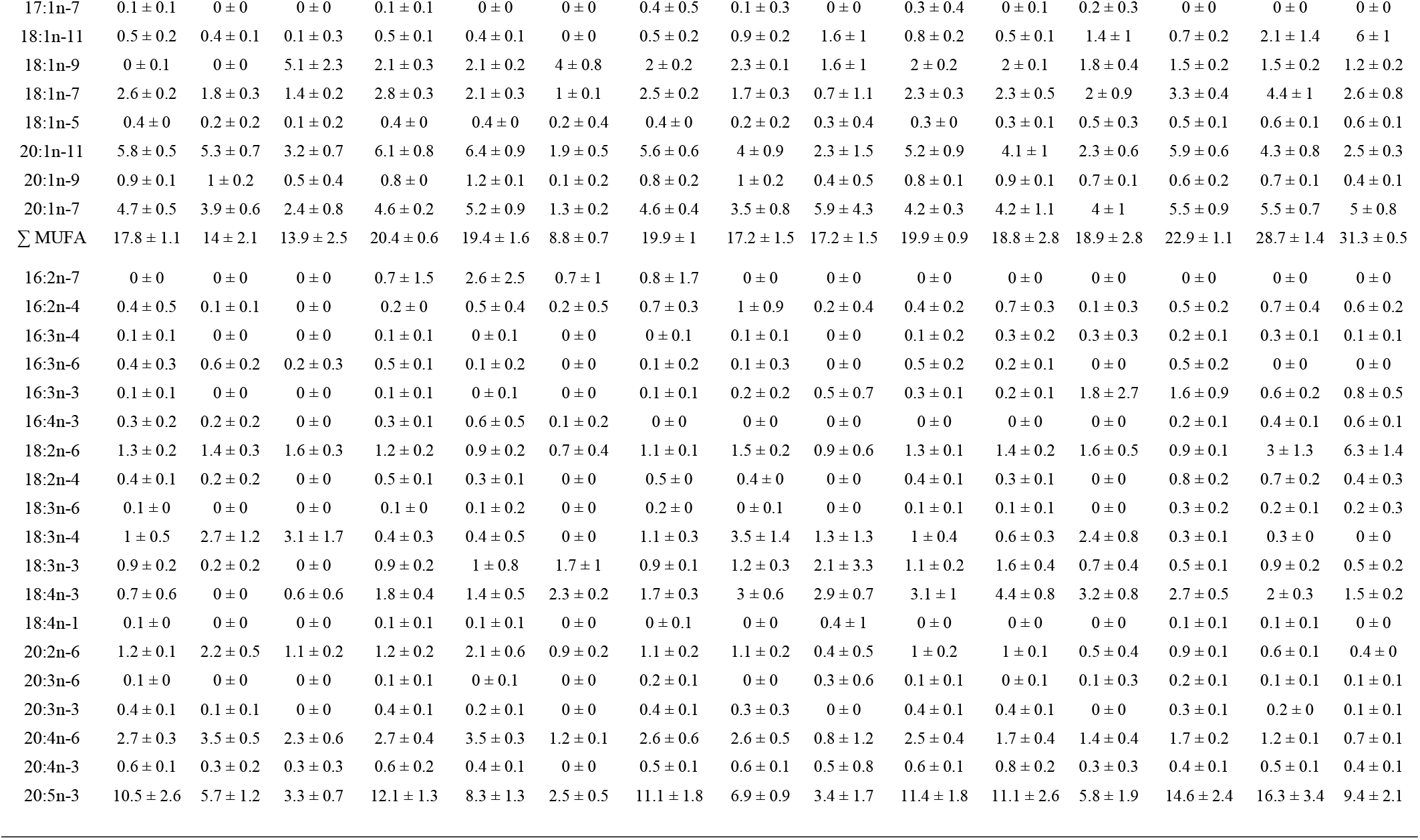
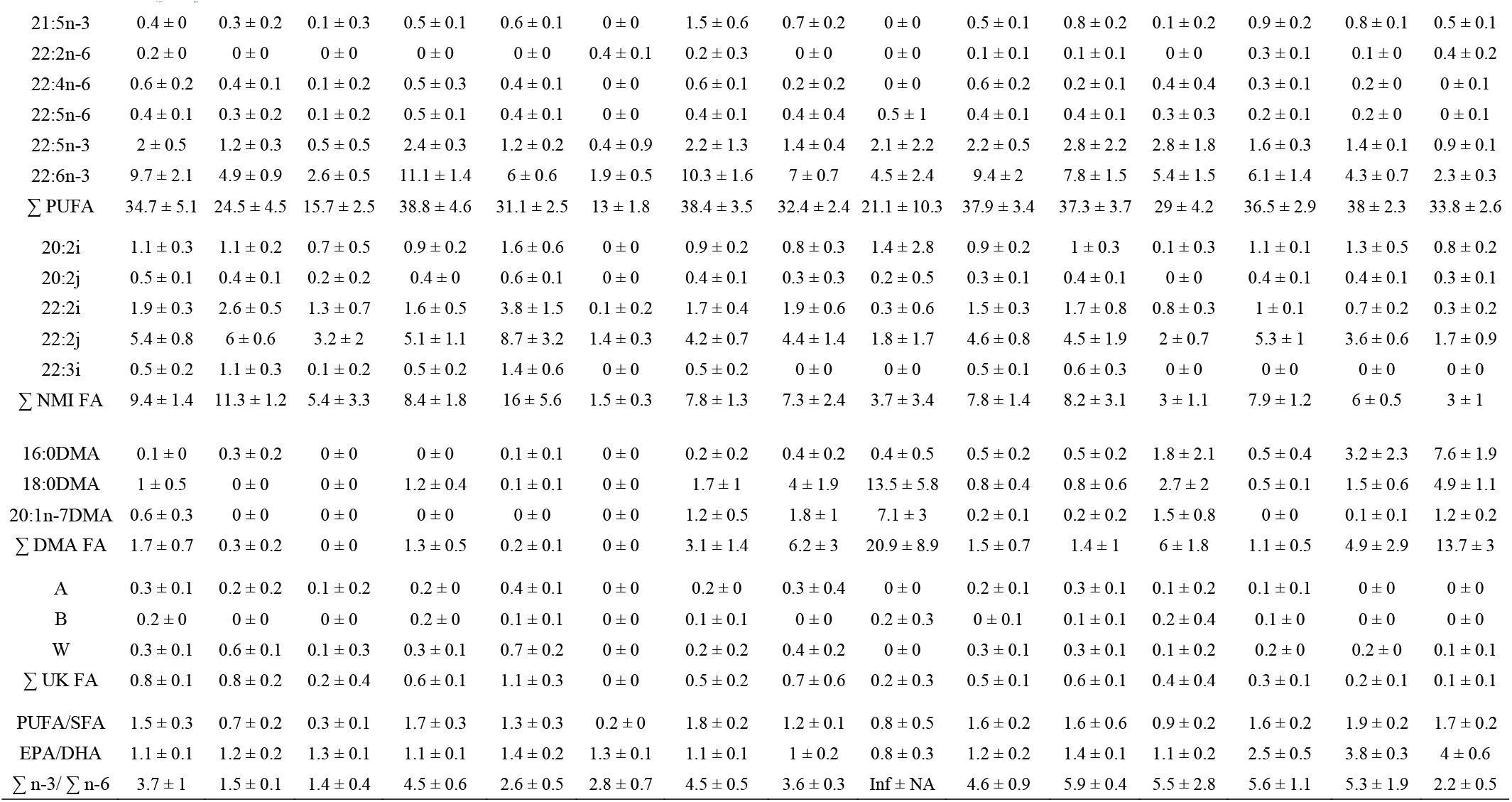
Fatty acid (FA) (%, mean ± SD, n = 5) composition of the three ontogenic stages of *Crepidula fornicata* (motile males, sessile males, sessile females) over the sampling survey. Only FA accounting for more than 0.5 % of total FA in at least one sample was shown. BFA: Branched FA; SFA: saturated FA; MUFA: monounsaturated FA; PUFA: polyunsaturated FA; NMI FA: non-methyl-interrupted FA; DMA: Dimethyl acetals FA; EPA: 20:5n-3; DHA: 22:6n-3.

| Fatty acids | S1 = 26<sup>th</sup> February |  |  | S2 = 21<sup>th</sup> March |  |  | S3 = 28<sup>th</sup> March |  |  | S4 = 12<sup>th</sup> April |  |  | S5 = 14<sup>th</sup> June |  |  |
| --- | --- | --- | --- | --- | --- | --- | --- | --- | --- | --- | --- | --- | --- | --- | --- |
|  | Sessile female | Sessile male | Motile male | Sessile female | Sessile male | Motile male | Sessile female | Sessile male | Motile male | Sessile female | Sessile male | Motile male | Sessile female | Sessile male | Motile male |
| TMTD | 1.3 ± 0.3 | 2.1 ± 1.1 | 0.8 ± 0.6 | 1.3 ± 0.2 | 0 ± 0.1 | 0.5 ± 0.4 | 1.3 ± 0.7 | 2.5 ± 0.4 | 0.9 ± 1.2 | 2.1 ± 1 | 2.9 ± 0.8 | 2.5 ± 1 | 2.1 ± 1.7 | 2 ± 0.7 | 1 ± 0.6 |
| 15:0iso | 0.2 ± 0 | 0 ± 0 | 0 ± 0 | 0.1 ± 0.1 | 0.1 ± 0.1 | 0.1 ± 0.2 | 0.1 ± 0.1 | 0 ± 0 | 0 ± 0 | 0.1 ± 0.1 | 0.1 ± 0.1 | 0.3 ± 0.4 | 0.1 ± 0.1 | 0.5 ± 0.3 | 1 ± 0.2 |
| 15:0ant | 0.1 ± 0 | 0 ± 0 | 0 ± 0 | 0 ± 0 | 0 ± 0 | 0 ± 0 | 0 ± 0.1 | 0 ± 0 | 0 ± 0 | 0.1 ± 0.1 | 0 ± 0 | 0.3 ± 0.5 | 0 ± 0 | 0.4 ± 0.3 | 0.7 ± 0.1 |
| 16:0iso | 1.3 ± 0.5 | 1.1 ± 0.3 | 1.1 ± 0.1 | 0.7 ± 0.1 | 0.9 ± 0.2 | 0.1 ± 0.2 | 0.9 ± 0.1 | 1 ± 0.1 | 0.6 ± 1 | 0.8 ± 0.2 | 0.5 ± 0.1 | 0.6 ± 0.4 | 0.7 ± 0.2 | 0.3 ± 0 | 0.4 ± 0.2 |
| 17:0iso | 5.4 ± 1.8 | 4.1 ± 1 | 3.4 ± 0.5 | 3.1 ± 1.8 | 1.7 ± 2.4 | 1.1 ± 1 | 3.5 ± 2 | 2.8 ± 1.4 | 2.5 ± 1.1 | 3.8 ± 1 | 2.4 ± 0.1 | 3 ± 0.3 | 3 ± 1 | 1.3 ± 0.2 | 1.4 ± 0.3 |
| 17:0ant | 3.6 ± 1.6 | 3.2 ± 0.7 | 2.4 ± 0.6 | 2.1 ± 0.5 | 3.5 ± 1 | 1.5 ± 0.3 | 2.8 ± 0.7 | 3.1 ± 0.4 | 3 ± 1 | 2.4 ± 0.7 | 1.8 ± 0.2 | 2.5 ± 0.3 | 2.4 ± 0.9 | 1.2 ± 0.2 | 1.5 ± 0.3 |
| 18:0iso | 0.7 ± 0.4 | 0.6 ± 0.4 | 0.3 ± 0.4 | 0.3 ± 0.3 | 0.2 ± 0.3 | 0 ± 0 | 0.5 ± 0.5 | 0.9 ± 0.2 | 0.2 ± 0.4 | 0.4 ± 0.6 | 0.7 ± 0.1 | 2.1 ± 2.9 | 0.5 ± 0.4 | 0.1 ± 0.1 | 0 ± 0 |
| $\Sigma$ BFA | 11.2 ± 4.3 | 8.9 ± 2.1 | 7.1 ± 1.3 | 6.3 ± 2.3 | 6.4 ± 2.6 | 2.8 ± 1 | 7.8 ± 2.6 | 7.8 ± 1.8 | 6.3 ± 2.2 | 7.5 ± 2.2 | 5.6 ± 0.3 | 8.9 ± 4 | 6.6 ± 2.6 | 3.8 ± 1 | 5 ± 0.9 |
| 14:0 | 1.5 ± 0.3 | 1.3 ± 0.7 | 1.4 ± 0.4 | 1.3 ± 0.2 | 3 ± 0.9 | 1.6 ± 0.1 | 1.3 ± 0.4 | 1.4 ± 0.3 | 0.8 ± 0.8 | 2 ± 0.5 | 2.3 ± 0.4 | 1.9 ± 0.7 | 2.4 ± 0.6 | 2.7 ± 0.6 | 2.2 ± 0.2 |
| 15:0 | 0.6 ± 0.1 | 0.6 ± 0.2 | 0.8 ± 0.2 | 0.4 ± 0 | 0.5 ± 0.1 | 0.6 ± 0.4 | 0.5 ± 0.1 | 0.4 ± 0 | 0.1 ± 0.3 | 0.5 ± 0.1 | 0.5 ± 0.1 | 0.6 ± 0 | 0.4 ± 0 | 0.7 ± 0.4 | 1.5 ± 0.3 |
| 16:0 | 11.9 ± 0.8 | 16.7 ± 3.1 | 25.2 ± 3.5 | 11.7 ± 1 | 11.6 ± 3.3 | 33.7 ± 1.7 | 10.9 ± 1 | 13 ± 0.9 | 12.3 ± 2.3 | 12 ± 0.4 | 12.6 ± 3.1 | 14.2 ± 2.1 | 11.2 ± 0.9 | 9.7 ± 1.4 | 7.6 ± 0.9 |
| 17:0 | 1.5 ± 0.1 | 1.2 ± 0.3 | 1 ± 0.1 | 1.4 ± 0.1 | 1.2 ± 0.1 | 0.7 ± 0.4 | 1 ± 0.5 | 0.8 ± 0.1 | 0.4 ± 0.5 | 0.9 ± 0.5 | 0.8 ± 0.1 | 1 ± 0.1 | 1.1 ± 0.2 | 0.7 ± 0.1 | 0.7 ± 0.1 |
| 18:0 | 6.3 ± 0.9 | 17.7 ± 7.3 | 27.5 ± 4.2 | 6.8 ± 0.7 | 9.2 ± 3.9 | 35.6 ± 0.9 | 6.6 ± 0.9 | 10.4 ± 1.3 | 14.8 ± 4.5 | 6.9 ± 0.4 | 9.1 ± 5.6 | 14.3 ± 3.1 | 6.9 ± 1.2 | 5.6 ± 1.1 | 7.5 ± 1.1 |
| 20:0 | 0.5 ± 0.1 | 0.7 ± 0.2 | 0.5 ± 0.3 | 0.5 ± 0.3 | 0.2 ± 0.2 | 0.6 ± 0.3 | 0.2 ± 0 | 0.2 ± 0.3 | 0 ± 0 | 0.2 ± 0 | 0.2 ± 0.1 | 0.1 ± 0.2 | 0.4 ± 0.1 | 0.2 ± 0.1 | 0.3 ± 0 |
| 22:0 | 0.5 ± 0.1 | 0.3 ± 0.2 | 0.1 ± 0.2 | 0.4 ± 0.1 | 0.2 ± 0 | 0 ± 0 | 0.4 ± 0.1 | 0.1 ± 0.1 | 0 ± 0 | 0.3 ± 0.1 | 0.2 ± 0.1 | 0 ± 0 | 0.3 ± 0.1 | 0 ± 0 | 0 ± 0 |
| 24:0 | 0.4 ± 0.1 | 0.6 ± 0.4 | 0.6 ± 0.4 | 0.2 ± 0 | 0.2 ± 0.1 | 0.7 ± 0.2 | 0.3 ± 0.2 | 0.1 ± 0.1 | 1.6 ± 1.8 | 0.4 ± 0.1 | 0.3 ± 0.1 | 1.2 ± 0.5 | 0.3 ± 0.1 | 0.1 ± 0.1 | 0 ± 0 |
| $\Sigma$ SFA | 23.2 ± 1.5 | 39.2 ± 9.6 | 57 ± 8.2 | 22.9 ± 1.5 | 26 ± 7.1 | 73.5 ± 2.4 | 21.4 ± 1.1 | 26.4 ± 1.7 | 30.1 ± 5.9 | 23.3 ± 0.9 | 25.8 ± 8.6 | 33.3 ± 4.6 | 23.1 ± 1.7 | 19.9 ± 1.2 | 19.7 ± 1.6 |
| 14:1n-5 | 0 ± 0 | 0 ± 0 | 0 ± 0 | 0 ± 0 | 0 ± 0 | 0 ± 0 | 0.1 ± 0.1 | 0 ± 0 | 0 ± 0 | 0 ± 0 | 0.1 ± 0.1 | 0.2 ± 0.5 | 0.1 ± 0.1 | 0 ± 0 | 0 ± 0 |
| 16:1n-11 | 0 ± 0 | 0 ± 0 | 0 ± 0 | 0 ± 0 | 0 ± 0 | 0 ± 0 | 0.4 ± 0.1 | 0.7 ± 0.4 | 0 ± 0 | 0.1 ± 0.1 | 0.2 ± 0.1 | 1.1 ± 1 | 0 ± 0 | 0.1 ± 0.1 | 0.2 ± 0.1 |
| 16:1n-9 | 0.7 ± 0.2 | 0.5 ± 0.3 | 0.3 ± 0.3 | 0.7 ± 0.2 | 0.1 ± 0.2 | 0.1 ± 0.2 | 0.5 ± 0.1 | 0.7 ± 0.5 | 0.4 ± 1 | 0.5 ± 0 | 0.6 ± 0.1 | 0.7 ± 0.3 | 0.3 ± 0 | 0.5 ± 0.4 | 4.1 ± 2.9 |
| 16:1n-7 | 1.4 ± 0.6 | 0.9 ± 0.2 | 0.8 ± 0.3 | 1.7 ± 0.2 | 1.3 ± 0.4 | 0.3 ± 0.4 | 1.5 ± 0.3 | 1.2 ± 0.4 | 2.6 ± 1.3 | 2.3 ± 0.6 | 2.9 ± 0.8 | 2.1 ± 0.3 | 3.6 ± 0.9 | 5.8 ± 1.9 | 6.4 ± 1.9 |
| 16:1n-5 | 0.4 ± 0.4 | 0 ± 0 | 0 ± 0 | 0.3 ± 0.1 | 0.1 ± 0.1 | 0 ± 0 | 0.3 ± 0.1 | 0.8 ± 0.4 | 1.4 ± 1.4 | 0.5 ± 0.2 | 0.4 ± 0.2 | 1.8 ± 1.2 | 0.9 ± 0.4 | 3.4 ± 2.1 | 2.2 ± 4.3 |
| 17:1n-7 | 0.1 ± 0.1 | 0 ± 0 | 0 ± 0 | 0.1 ± 0.1 | 0 ± 0 | 0 ± 0 | 0.4 ± 0.5 | 0.1 ± 0.3 | 0 ± 0 | 0.3 ± 0.4 | 0 ± 0.1 | 0.2 ± 0.3 | 0 ± 0 | 0 ± 0 | 0 ± 0 |
| 18:1n-11 | 0.5 ± 0.2 | 0.4 ± 0.1 | 0.1 ± 0.3 | 0.5 ± 0.1 | 0.4 ± 0.1 | 0 ± 0 | 0.5 ± 0.2 | 0.9 ± 0.2 | 1.6 ± 1 | 0.8 ± 0.2 | 0.5 ± 0.1 | 1.4 ± 1 | 0.7 ± 0.2 | 2.1 ± 1.4 | 6 ± 1 |
| 18:1n-9 | 0 ± 0.1 | 0 ± 0 | 5.1 ± 2.3 | 2.1 ± 0.3 | 2.1 ± 0.2 | 4 ± 0.8 | 2 ± 0.2 | 2.3 ± 0.1 | 1.6 ± 1 | 2 ± 0.2 | 2 ± 0.1 | 1.8 ± 0.4 | 1.5 ± 0.2 | 1.5 ± 0.2 | 1.2 ± 0.2 |
| 18:1n-7 | 2.6 ± 0.2 | 1.8 ± 0.3 | 1.4 ± 0.2 | 2.8 ± 0.3 | 2.1 ± 0.3 | 1 ± 0.1 | 2.5 ± 0.2 | 1.7 ± 0.3 | 0.7 ± 1.1 | 2.3 ± 0.3 | 2.3 ± 0.5 | 2 ± 0.9 | 3.3 ± 0.4 | 4.4 ± 1 | 2.6 ± 0.8 |
| 18:1n-5 | 0.4 ± 0 | 0.2 ± 0.2 | 0.1 ± 0.2 | 0.4 ± 0 | 0.4 ± 0 | 0.2 ± 0.4 | 0.4 ± 0 | 0.2 ± 0.2 | 0.3 ± 0.4 | 0.3 ± 0 | 0.3 ± 0.1 | 0.5 ± 0.3 | 0.5 ± 0.1 | 0.6 ± 0.1 | 0.6 ± 0.1 |
| 20:1n-11 | 5.8 ± 0.5 | 5.3 ± 0.7 | 3.2 ± 0.7 | 6.1 ± 0.8 | 6.4 ± 0.9 | 1.9 ± 0.5 | 5.6 ± 0.6 | 4 ± 0.9 | 2.3 ± 1.5 | 5.2 ± 0.9 | 4.1 ± 1 | 2.3 ± 0.6 | 5.9 ± 0.6 | 4.3 ± 0.8 | 2.5 ± 0.3 |
| 20:1n-9 | 0.9 ± 0.1 | 1 ± 0.2 | 0.5 ± 0.4 | 0.8 ± 0 | 1.2 ± 0.1 | 0.1 ± 0.2 | 0.8 ± 0.2 | 1 ± 0.2 | 0.4 ± 0.5 | 0.8 ± 0.1 | 0.9 ± 0.1 | 0.7 ± 0.1 | 0.6 ± 0.2 | 0.7 ± 0.1 | 0.4 ± 0.1 |
| 20:1n-7 | 4.7 ± 0.5 | 3.9 ± 0.6 | 2.4 ± 0.8 | 4.6 ± 0.2 | 5.2 ± 0.9 | 1.3 ± 0.2 | 4.6 ± 0.4 | 3.5 ± 0.8 | 5.9 ± 4.3 | 4.2 ± 0.3 | 4.2 ± 1.1 | 4 ± 1 | 5.5 ± 0.9 | 5.5 ± 0.7 | 5 ± 0.8 |
| Σ MUFA | 17.8 ± 1.1 | 14 ± 2.1 | 13.9 ± 2.5 | 20.4 ± 0.6 | 19.4 ± 1.6 | 8.8 ± 0.7 | 19.9 ± 1 | 17.2 ± 1.5 | 17.2 ± 1.5 | 19.9 ± 0.9 | 18.8 ± 2.8 | 18.9 ± 2.8 | 22.9 ± 1.1 | 28.7 ± 1.4 | 31.3 ± 0.5 |
| 16:2n-7 | 0 ± 0 | 0 ± 0 | 0 ± 0 | 0.7 ± 1.5 | 2.6 ± 2.5 | 0.7 ± 1 | 0.8 ± 1.7 | 0 ± 0 | 0 ± 0 | 0 ± 0 | 0 ± 0 | 0 ± 0 | 0 ± 0 | 0 ± 0 | 0 ± 0 |
| 16:2n-4 | 0.4 ± 0.5 | 0.1 ± 0.1 | 0 ± 0 | 0.2 ± 0 | 0.5 ± 0.4 | 0.2 ± 0.5 | 0.7 ± 0.3 | 1 ± 0.9 | 0.2 ± 0.4 | 0.4 ± 0.2 | 0.7 ± 0.3 | 0.1 ± 0.3 | 0.5 ± 0.2 | 0.7 ± 0.4 | 0.6 ± 0.2 |
| 16:3n-4 | 0.1 ± 0.1 | 0 ± 0 | 0 ± 0 | 0.1 ± 0.1 | 0 ± 0.1 | 0 ± 0 | 0 ± 0.1 | 0.1 ± 0.1 | 0 ± 0 | 0.1 ± 0.2 | 0.3 ± 0.2 | 0.3 ± 0.3 | 0.2 ± 0.1 | 0.3 ± 0.1 | 0.1 ± 0.1 |
| 16:3n-6 | 0.4 ± 0.3 | 0.6 ± 0.2 | 0.2 ± 0.3 | 0.5 ± 0.1 | 0.1 ± 0.2 | 0 ± 0 | 0.1 ± 0.2 | 0.1 ± 0.3 | 0 ± 0 | 0.5 ± 0.2 | 0.2 ± 0.1 | 0 ± 0 | 0.5 ± 0.2 | 0 ± 0 | 0 ± 0 |
| 16:3n-3 | 0.1 ± 0.1 | 0 ± 0 | 0 ± 0 | 0.1 ± 0.1 | 0 ± 0.1 | 0 ± 0 | 0.1 ± 0.1 | 0.2 ± 0.2 | 0.5 ± 0.7 | 0.3 ± 0.1 | 0.2 ± 0.1 | 1.8 ± 2.7 | 1.6 ± 0.9 | 0.6 ± 0.2 | 0.8 ± 0.5 |
| 16:4n-3 | 0.3 ± 0.2 | 0.2 ± 0.2 | 0 ± 0 | 0.3 ± 0.1 | 0.6 ± 0.5 | 0.1 ± 0.2 | 0 ± 0 | 0 ± 0 | 0 ± 0 | 0 ± 0 | 0 ± 0 | 0 ± 0 | 0.2 ± 0.1 | 0.4 ± 0.1 | 0.6 ± 0.1 |
| 18:2n-6 | 1.3 ± 0.2 | 1.4 ± 0.3 | 1.6 ± 0.3 | 1.2 ± 0.2 | 0.9 ± 0.2 | 0.7 ± 0.4 | 1.1 ± 0.1 | 1.5 ± 0.2 | 0.9 ± 0.6 | 1.3 ± 0.1 | 1.4 ± 0.2 | 1.6 ± 0.5 | 0.9 ± 0.1 | 3 ± 1.3 | 6.3 ± 1.4 |
| 18:2n-4 | 0.4 ± 0.1 | 0.2 ± 0.2 | 0 ± 0 | 0.5 ± 0.1 | 0.3 ± 0.1 | 0 ± 0 | 0.5 ± 0 | 0.4 ± 0 | 0 ± 0 | 0.4 ± 0.1 | 0.3 ± 0.1 | 0 ± 0 | 0.8 ± 0.2 | 0.7 ± 0.2 | 0.4 ± 0.3 |
| 18:3n-6 | 0.1 ± 0 | 0 ± 0 | 0 ± 0 | 0.1 ± 0 | 0.1 ± 0.2 | 0 ± 0 | 0.2 ± 0 | 0 ± 0.1 | 0 ± 0 | 0.1 ± 0.1 | 0.1 ± 0.1 | 0 ± 0 | 0.3 ± 0.2 | 0.2 ± 0.1 | 0.2 ± 0.3 |
| 18:3n-4 | 1 ± 0.5 | 2.7 ± 1.2 | 3.1 ± 1.7 | 0.4 ± 0.3 | 0.4 ± 0.5 | 0 ± 0 | 1.1 ± 0.3 | 3.5 ± 1.4 | 1.3 ± 1.3 | 1 ± 0.4 | 0.6 ± 0.3 | 2.4 ± 0.8 | 0.3 ± 0.1 | 0.3 ± 0 | 0 ± 0 |
| 18:3n-3 | 0.9 ± 0.2 | 0.2 ± 0.2 | 0 ± 0 | 0.9 ± 0.2 | 1 ± 0.8 | 1.7 ± 1 | 0.9 ± 0.1 | 1.2 ± 0.3 | 2.1 ± 3.3 | 1.1 ± 0.2 | 1.6 ± 0.4 | 0.7 ± 0.4 | 0.5 ± 0.1 | 0.9 ± 0.2 | 0.5 ± 0.2 |
| 18:4n-3 | 0.7 ± 0.6 | 0 ± 0 | 0.6 ± 0.6 | 1.8 ± 0.4 | 1.4 ± 0.5 | 2.3 ± 0.2 | 1.7 ± 0.3 | 3 ± 0.6 | 2.9 ± 0.7 | 3.1 ± 1 | 4.4 ± 0.8 | 3.2 ± 0.8 | 2.7 ± 0.5 | 2 ± 0.3 | 1.5 ± 0.2 |
| 18:4n-1 | 0.1 ± 0 | 0 ± 0 | 0 ± 0 | 0.1 ± 0.1 | 0.1 ± 0.1 | 0 ± 0 | 0 ± 0.1 | 0 ± 0 | 0.4 ± 1 | 0 ± 0 | 0 ± 0 | 0 ± 0 | 0.1 ± 0.1 | 0.1 ± 0.1 | 0 ± 0 |
| 20:2n-6 | 1.2 ± 0.1 | 2.2 ± 0.5 | 1.1 ± 0.2 | 1.2 ± 0.2 | 2.1 ± 0.6 | 0.9 ± 0.2 | 1.1 ± 0.2 | 1.1 ± 0.2 | 0.4 ± 0.5 | 1 ± 0.2 | 1 ± 0.1 | 0.5 ± 0.4 | 0.9 ± 0.1 | 0.6 ± 0.1 | 0.4 ± 0 |
| 20:3n-6 | 0.1 ± 0 | 0 ± 0 | 0 ± 0 | 0.1 ± 0.1 | 0 ± 0.1 | 0 ± 0 | 0.2 ± 0.1 | 0 ± 0 | 0.3 ± 0.6 | 0.1 ± 0.1 | 0 ± 0.1 | 0.1 ± 0.3 | 0.2 ± 0.1 | 0.1 ± 0.1 | 0.1 ± 0.1 |
| 20:3n-3 | 0.4 ± 0.1 | 0.1 ± 0.1 | 0 ± 0 | 0.4 ± 0.1 | 0.2 ± 0.1 | 0 ± 0 | 0.4 ± 0.1 | 0.3 ± 0.3 | 0 ± 0 | 0.4 ± 0.1 | 0.4 ± 0.1 | 0 ± 0 | 0.3 ± 0.1 | 0.2 ± 0 | 0.1 ± 0.1 |
| 20:4n-6 | 2.7 ± 0.3 | 3.5 ± 0.5 | 2.3 ± 0.6 | 2.7 ± 0.4 | 3.5 ± 0.3 | 1.2 ± 0.1 | 2.6 ± 0.6 | 2.6 ± 0.5 | 0.8 ± 1.2 | 2.5 ± 0.4 | 1.7 ± 0.4 | 1.4 ± 0.4 | 1.7 ± 0.2 | 1.2 ± 0.1 | 0.7 ± 0.1 |
| 20:4n-3 | 0.6 ± 0.1 | 0.3 ± 0.2 | 0.3 ± 0.3 | 0.6 ± 0.2 | 0.4 ± 0.1 | 0 ± 0 | 0.5 ± 0.1 | 0.6 ± 0.1 | 0.5 ± 0.8 | 0.6 ± 0.1 | 0.8 ± 0.2 | 0.3 ± 0.3 | 0.4 ± 0.1 | 0.5 ± 0.1 | 0.4 ± 0.1 |
| 20:5n-3 | 10.5 ± 2.6 | 5.7 ± 1.2 | 3.3 ± 0.7 | 12.1 ± 1.3 | 8.3 ± 1.3 | 2.5 ± 0.5 | 11.1 ± 1.8 | 6.9 ± 0.9 | 3.4 ± 1.7 | 11.4 ± 1.8 | 11.1 ± 2.6 | 5.8 ± 1.9 | 14.6 ± 2.4 | 16.3 ± 3.4 | 9.4 ± 2.1 |
| 21:5n-3 | 0.4 ± 0 | 0.3 ± 0.2 | 0.1 ± 0.3 | 0.5 ± 0.1 | 0.6 ± 0.1 | 0 ± 0 | 1.5 ± 0.6 | 0.7 ± 0.2 | 0 ± 0 | 0.5 ± 0.1 | 0.8 ± 0.2 | 0.1 ± 0.2 | 0.9 ± 0.2 | 0.8 ± 0.1 | 0.5 ± 0.1 |
| 22:2n-6 | 0.2 ± 0 | 0 ± 0 | 0 ± 0 | 0 ± 0 | 0 ± 0 | 0.4 ± 0.1 | 0.2 ± 0.3 | 0 ± 0 | 0 ± 0 | 0.1 ± 0.1 | 0.1 ± 0.1 | 0 ± 0 | 0.3 ± 0.1 | 0.1 ± 0 | 0.4 ± 0.2 |
| 22:4n-6 | 0.6 ± 0.2 | 0.4 ± 0.1 | 0.1 ± 0.2 | 0.5 ± 0.3 | 0.4 ± 0.1 | 0 ± 0 | 0.6 ± 0.1 | 0.2 ± 0.2 | 0 ± 0 | 0.6 ± 0.2 | 0.2 ± 0.1 | 0.4 ± 0.4 | 0.3 ± 0.1 | 0.2 ± 0 | 0 ± 0.1 |
| 22:5n-6 | 0.4 ± 0.1 | 0.3 ± 0.2 | 0.1 ± 0.2 | 0.5 ± 0.1 | 0.4 ± 0.1 | 0 ± 0 | 0.4 ± 0.1 | 0.4 ± 0.4 | 0.5 ± 1 | 0.4 ± 0.1 | 0.4 ± 0.1 | 0.3 ± 0.3 | 0.2 ± 0.1 | 0.2 ± 0 | 0 ± 0.1 |
| 22:5n-3 | 2 ± 0.5 | 1.2 ± 0.3 | 0.5 ± 0.5 | 2.4 ± 0.3 | 1.2 ± 0.2 | 0.4 ± 0.9 | 2.2 ± 1.3 | 1.4 ± 0.4 | 2.1 ± 2.2 | 2.2 ± 0.5 | 2.8 ± 2.2 | 2.8 ± 1.8 | 1.6 ± 0.3 | 1.4 ± 0.1 | 0.9 ± 0.1 |
| 22:6n-3 | 9.7 ± 2.1 | 4.9 ± 0.9 | 2.6 ± 0.5 | 11.1 ± 1.4 | 6 ± 0.6 | 1.9 ± 0.5 | 10.3 ± 1.6 | 7 ± 0.7 | 4.5 ± 2.4 | 9.4 ± 2 | 7.8 ± 1.5 | 5.4 ± 1.5 | 6.1 ± 1.4 | 4.3 ± 0.7 | 2.3 ± 0.3 |
| Σ PUFA | 34.7 ± 5.1 | 24.5 ± 4.5 | 15.7 ± 2.5 | 38.8 ± 4.6 | 31.1 ± 2.5 | 13 ± 1.8 | 38.4 ± 3.5 | 32.4 ± 2.4 | 21.1 ± 10.3 | 37.9 ± 3.4 | 37.3 ± 3.7 | 29 ± 4.2 | 36.5 ± 2.9 | 38 ± 2.3 | 33.8 ± 2.6 |
| 20:2i | 1.1 ± 0.3 | 1.1 ± 0.2 | 0.7 ± 0.5 | 0.9 ± 0.2 | 1.6 ± 0.6 | 0 ± 0 | 0.9 ± 0.2 | 0.8 ± 0.3 | 1.4 ± 2.8 | 0.9 ± 0.2 | 1 ± 0.3 | 0.1 ± 0.3 | 1.1 ± 0.1 | 1.3 ± 0.5 | 0.8 ± 0.2 |
| 20:2j | 0.5 ± 0.1 | 0.4 ± 0.1 | 0.2 ± 0.2 | 0.4 ± 0 | 0.6 ± 0.1 | 0 ± 0 | 0.4 ± 0.1 | 0.3 ± 0.3 | 0.2 ± 0.5 | 0.3 ± 0.1 | 0.4 ± 0.1 | 0 ± 0 | 0.4 ± 0.1 | 0.4 ± 0.1 | 0.3 ± 0.1 |
| 22:2i | 1.9 ± 0.3 | 2.6 ± 0.5 | 1.3 ± 0.7 | 1.6 ± 0.5 | 3.8 ± 1.5 | 0.1 ± 0.2 | 1.7 ± 0.4 | 1.9 ± 0.6 | 0.3 ± 0.6 | 1.5 ± 0.3 | 1.7 ± 0.8 | 0.8 ± 0.3 | 1 ± 0.1 | 0.7 ± 0.2 | 0.3 ± 0.2 |
| 22:2j | 5.4 ± 0.8 | 6 ± 0.6 | 3.2 ± 2 | 5.1 ± 1.1 | 8.7 ± 3.2 | 1.4 ± 0.3 | 4.2 ± 0.7 | 4.4 ± 1.4 | 1.8 ± 1.7 | 4.6 ± 0.8 | 4.5 ± 1.9 | 2 ± 0.7 | 5.3 ± 1 | 3.6 ± 0.6 | 1.7 ± 0.9 |
| 22:3i | 0.5 ± 0.2 | 1.1 ± 0.3 | 0.1 ± 0.2 | 0.5 ± 0.2 | 1.4 ± 0.6 | 0 ± 0 | 0.5 ± 0.2 | 0 ± 0 | 0 ± 0 | 0.5 ± 0.1 | 0.6 ± 0.3 | 0 ± 0 | 0 ± 0 | 0 ± 0 | 0 ± 0 |
| Σ NMI FA | 9.4 ± 1.4 | 11.3 ± 1.2 | 5.4 ± 3.3 | 8.4 ± 1.8 | 16 ± 5.6 | 1.5 ± 0.3 | 7.8 ± 1.3 | 7.3 ± 2.4 | 3.7 ± 3.4 | 7.8 ± 1.4 | 8.2 ± 3.1 | 3 ± 1.1 | 7.9 ± 1.2 | 6 ± 0.5 | 3 ± 1 |
| 16:0DMA | 0.1 ± 0 | 0.3 ± 0.2 | 0 ± 0 | 0 ± 0 | 0.1 ± 0.1 | 0 ± 0 | 0.2 ± 0.2 | 0.4 ± 0.2 | 0.4 ± 0.5 | 0.5 ± 0.2 | 0.5 ± 0.2 | 1.8 ± 2.1 | 0.5 ± 0.4 | 3.2 ± 2.3 | 7.6 ± 1.9 |
| 18:0DMA | 1 ± 0.5 | 0 ± 0 | 0 ± 0 | 1.2 ± 0.4 | 0.1 ± 0.1 | 0 ± 0 | 1.7 ± 1 | 4 ± 1.9 | 13.5 ± 5.8 | 0.8 ± 0.4 | 0.8 ± 0.6 | 2.7 ± 2 | 0.5 ± 0.1 | 1.5 ± 0.6 | 4.9 ± 1.1 |
| 20:1n-7DMA | 0.6 ± 0.3 | 0 ± 0 | 0 ± 0 | 0 ± 0 | 0 ± 0 | 0 ± 0 | 1.2 ± 0.5 | 1.8 ± 1 | 7.1 ± 3 | 0.2 ± 0.1 | 0.2 ± 0.2 | 1.5 ± 0.8 | 0 ± 0 | 0.1 ± 0.1 | 1.2 ± 0.2 |
| Σ DMA FA | 1.7 ± 0.7 | 0.3 ± 0.2 | 0 ± 0 | 1.3 ± 0.5 | 0.2 ± 0.1 | 0 ± 0 | 3.1 ± 1.4 | 6.2 ± 3 | 20.9 ± 8.9 | 1.5 ± 0.7 | 1.4 ± 1 | 6 ± 1.8 | 1.1 ± 0.5 | 4.9 ± 2.9 | 13.7 ± 3 |
| A | 0.3 ± 0.1 | 0.2 ± 0.2 | 0.1 ± 0.2 | 0.2 ± 0 | 0.4 ± 0.1 | 0 ± 0 | 0.2 ± 0 | 0.3 ± 0.4 | 0 ± 0 | 0.2 ± 0.1 | 0.3 ± 0.1 | 0.1 ± 0.2 | 0.1 ± 0.1 | 0 ± 0 | 0 ± 0 |
| B | 0.2 ± 0 | 0 ± 0 | 0 ± 0 | 0.2 ± 0 | 0.1 ± 0.1 | 0 ± 0 | 0.1 ± 0.1 | 0 ± 0 | 0.2 ± 0.3 | 0 ± 0.1 | 0.1 ± 0.1 | 0.2 ± 0.4 | 0.1 ± 0 | 0 ± 0 | 0 ± 0 |
| W | 0.3 ± 0.1 | 0.6 ± 0.1 | 0.1 ± 0.3 | 0.3 ± 0.1 | 0.7 ± 0.2 | 0 ± 0 | 0.2 ± 0.2 | 0.4 ± 0.2 | 0 ± 0 | 0.3 ± 0.1 | 0.3 ± 0.1 | 0.1 ± 0.2 | 0.2 ± 0 | 0.2 ± 0 | 0.1 ± 0.1 |
| Σ UK FA | 0.8 ± 0.1 | 0.8 ± 0.2 | 0.2 ± 0.4 | 0.6 ± 0.1 | 1.1 ± 0.3 | 0 ± 0 | 0.5 ± 0.2 | 0.7 ± 0.6 | 0.2 ± 0.3 | 0.5 ± 0.1 | 0.6 ± 0.1 | 0.4 ± 0.4 | 0.3 ± 0.1 | 0.2 ± 0.1 | 0.1 ± 0.1 |
| PUFA/SFA | 1.5 ± 0.3 | 0.7 ± 0.2 | 0.3 ± 0.1 | 1.7 ± 0.3 | 1.3 ± 0.3 | 0.2 ± 0 | 1.8 ± 0.2 | 1.2 ± 0.1 | 0.8 ± 0.5 | 1.6 ± 0.2 | 1.6 ± 0.6 | 0.9 ± 0.2 | 1.6 ± 0.2 | 1.9 ± 0.2 | 1.7 ± 0.2 |
| EPA/DHA | 1.1 ± 0.1 | 1.2 ± 0.2 | 1.3 ± 0.1 | 1.1 ± 0.1 | 1.4 ± 0.2 | 1.3 ± 0.1 | 1.1 ± 0.1 | 1 ± 0.2 | 0.8 ± 0.3 | 1.2 ± 0.2 | 1.4 ± 0.1 | 1.1 ± 0.2 | 2.5 ± 0.5 | 3.8 ± 0.3 | 4 ± 0.6 |
| Σ n-3/Σ n-6 | 3.7 ± 1 | 1.5 ± 0.1 | 1.4 ± 0.4 | 4.5 ± 0.6 | 2.6 ± 0.5 | 2.8 ± 0.7 | 4.5 ± 0.5 | 3.6 ± 0.3 | Inf ± NA | 4.6 ± 0.9 | 5.9 ± 0.4 | 5.5 ± 2.8 | 5.6 ± 1.1 | 5.3 ± 1.9 | 2.2 ± 0.5 |

**Table S5:** Stable isotope (mean ± SD, n = 5) composition of the three ontogenic stages of *Crepidula fornicata* (motile males, sessile males, sessile females) over the sampling survey.

|  | S1 = 26<sup>th</sup> February |  |  | S2 = 21<sup>th</sup> March |  |  | S3 = 28<sup>th</sup> March |  |  | S4 = 12<sup>th</sup> April |  |  | S5 = 14<sup>th</sup> June |  |  |
| --- | --- | --- | --- | --- | --- | --- | --- | --- | --- | --- | --- | --- | --- | --- | --- |
|  | Sessile female | Sessile male | Motile male | Sessile female | Sessile male | Motile male | Sessile female | Sessile male | Motile male | Sessile female | Sessile male | Motile male | Sessile female | Sessile male | Motile male |
| $\delta^{13}\text{C}$ | -19.5 ± 0.3 | -19.4 ± 0.4 | -19.4 ± 0.5 | -19.8 ± 0.3 | -19.7 ± 0.3 | -20.1 ± 0.5 | -19.2 ± 0.6 | -19.6 ± 0.4 | -19.9 ± 0.3 | -20.1 ± 0.4 | -20.3 ± 0.8 | -20.6 ± 0.5 | -18.1 ± 0.3 | -18.5 ± 0.1 | -18.5 ± 0.3 |
| $\delta^{15}\text{N}$ | 9.3 ± 0.4 | 8.9 ± 0.8 | 8.9 ± 0.3 | 9.6 ± 0.5 | 8.9 ± 0.5 | 8.4 ± 1.1 | 9.7 ± 0.3 | 9.4 ± 0.6 | 8.5 ± 0.4 | 8.4 ± 0.7 | 8.4 ± 0.6 | 8.3 ± 0.5 | 8.6 ± 0.6 | 8.3 ± 0.5 | 7.5 ± 0.3 |
| C/N | 4 ± 0.3 | 3.8 ± 0.8 | 4 ± 0.2 | 3.2 ± 0.8 | 3.8 ± 0.6 | 3.5 ± 0.7 | 3.6 ± 0.4 | 3.3 ± 0.4 | 3.8 ± 0.2 | 3.6 ± 0.4 | 4.3 ± 0.2 | 3.8 ± 0.2 | 3.5 ± 0.6 | 4.1 ± 1.2 | 3.4 ± 0.3 |

**Figure S1:**
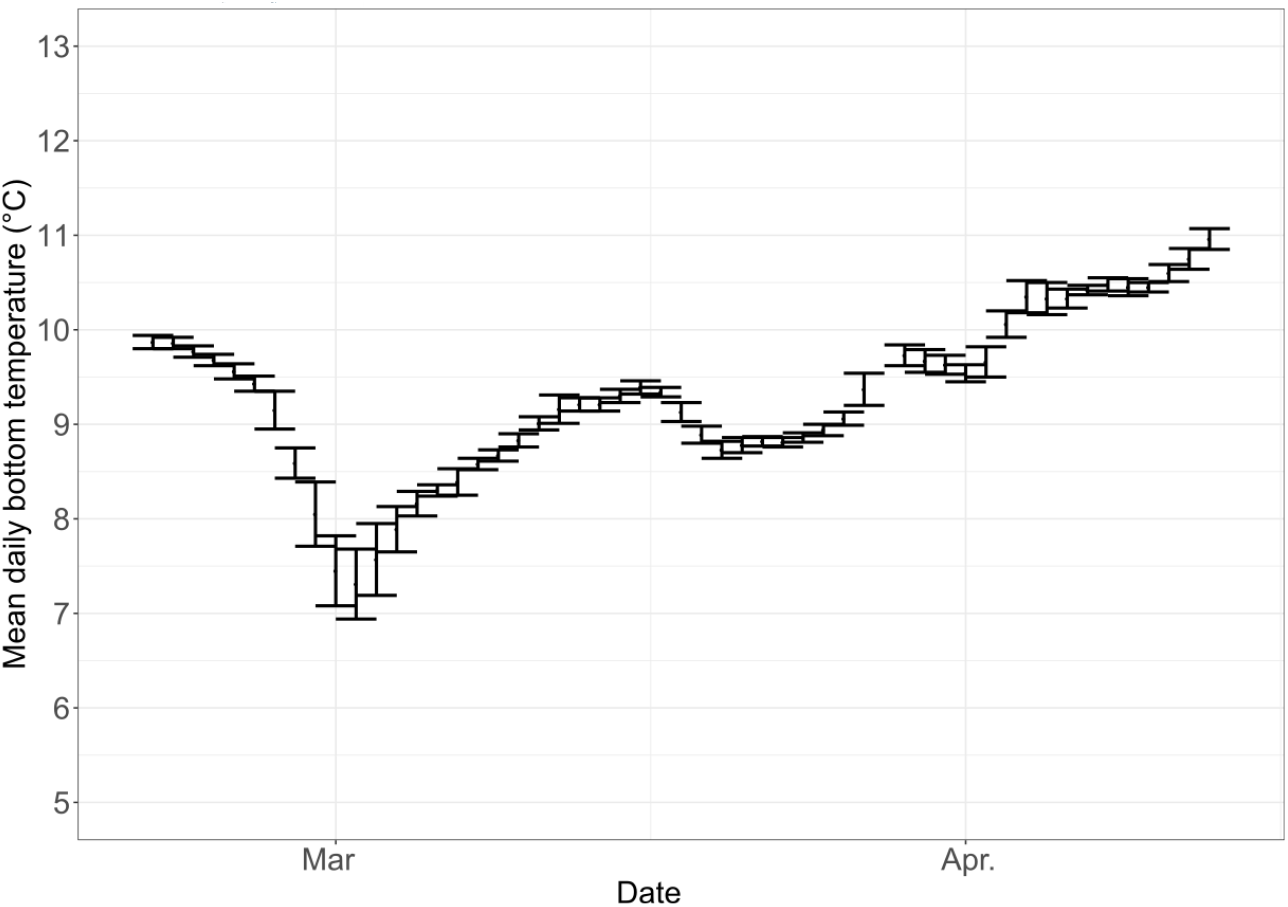
Mean daily temperature (± SD) at the bottom in our study site.

